# Antibody landscape of C57BL/6 mice cured of B78 melanoma via immunotherapy

**DOI:** 10.1101/2023.02.24.529012

**Authors:** A Hoefges, SJ McIlwain, AK Erbe, N Mathers, A Xu, D Melby, K Tetreault, T Le, K Kim, RS Pinapati, B Garcia, J Patel, M Heck, AS Feils, N Tsarovsky, JA Hank, ZS Morris, IM Ong, PM Sondel

**Author notes:** Corresponding Author: Paul M. Sondel, M.D., Ph.D. Reed and Carolee Walker, Professor of Childhood Cancer Research Departments of Pediatrics, Human Oncology and Genetics; 4159 MACC Fund Childhood Cancer Research Wing Wisconsin Institutes for Medical Research University of Wisconsin, Madison WI; – office; – cell. Co-Senior Authors (IM Ong, PM Sondel).

## Abstract

1

Antibodies can play an important role in innate and adaptive immune responses against cancer, and in preventing infectious disease. Flow cytometry analysis of sera of immune mice that were previously cured of their melanoma through a combined immunotherapy regimen with long-term memory showed strong antibody-binding against melanoma tumor cell lines. Using a high-density whole-proteome peptide array, we assessed potential protein-targets for antibodies found in immune sera. Sera from 6 of these cured mice were analyzed with this high-density, whole-proteome peptide array to determine specific antibody-binding sites and their linear peptide sequence. We identified thousands of peptides that were targeted by 2 or more of these 6 mice and exhibited strong antibody binding only by immune, not naive sera. Confirmatory studies were done to validate these results using 2 separate ELISA-based systems. To the best of our knowledge, this is the first study of the “immunome” of protein-based epitopes that are recognized by immune sera from mice cured of cancer via immunotherapy.

**summary:** Hoefges et al. utilized a whole-proteome peptide array approach to show that C57BL/6 mice develop a large repertoire of antibodies against linear peptide sequences of their melanoma after receiving a curative immunotherapy regimen consisting of radiation and an immunocytokine.

## 2 Introduction

Cancer immunotherapy has revolutionized cancer treatment and has helped thousands of patients (Couzin-Frankel, 2013; Patel & Minn, 2018). However, most patients are still not showing positive responses to current cancer immunotherapy treatment regimens (Chiriva-Internati & Bot, 2015; Patel & Minn, 2018). Using radiation therapy (RT) and intratumoral injections of immunocytokine (IC), we have developed a local *in-situ* vaccine (ISV, RT+IC) regimen capable of curing immunocompetent C57BL/6 mice bearing syngeneic B78 melanoma tumors and resulting in protective immune memory (Morris et al., 2016). Even though B78 is considered a functionally “cold” tumor due to its lack of response to checkpoint inhibitors (Gentles et al., 2015; Morris et al., 2018), our RT+IC regimen can cure many of them. With our *in-situ* vaccine, RT acts to increase the immunogenicity of the tumor by modifying its phenotype and releasing immune stimulatory cytokines. IC is an engineered fusion protein consisting of a tumor-specific monoclonal antibody targeting disialoganglioside (GD2) linked to IL2. GD2 is a molecule expressed on the surface of most neuroectodermal tumors and some nerve fibers. We also demonstrated that our *in-situ* vaccine causes epitope spread; 75% of cured mice reject a challenge with B16 melanoma cells (Morris et al., 2016; Yang et al., 2012). B16 melanoma cells do not express the GD2 antigen and are the parental cell line to B78 (Haraguchi et al., 1994; Silagi, 1969; Silagi et al., 1972). We observed strong antibody-binding to B16 cells using serum from cured as compared to naïve mice (Baniel et al., 2020). These antibodies might enable MHC-independent, CD8-T cell independent anti-tumor adaptive immune responses via macrophage-mediated antibody-dependent direct tumor cell killing (Jagodinsky et al., 2022). However, the exact antigen targets of these endogenous antibodies are unknown.

Identifying epitopes on tumor cells that are recognized by antibodies may help identify the immunodominant antigens of cold human tumors, which may help in overcoming immune resistance in these cancers (Sasaki et al., 2020; Shen et al., 2013; Tarp et al., 2007). With the RT+IC regimen, although we are targeting GD2, the memory response does not require GD2 (Morris et al., 2016). Knowledge of these additional antigenic targets may help to identify biomarkers of positive responses and identify potential new therapeutic targets.

In this paper, we utilized a high-density peptide array approach to probe every protein of the mouse proteome, broken down into 16-mer peptides in a 2 or 4 amino acid (aa) tiling approach, to identify antibody targets, using serum from cured mice vs. their matched naïve sample. This high-density peptide array technology has been used for several productive applications recently (Engmark et al., 2016; Haj et al., 2020; Lo et al., 2020; Lyamichev et al., 2017; Mishra et al., 2021; Shen et al., 2019). Using this approach, we identified many tumor antigens expressed by cold murine tumors in individual mice as well as some tumor antigens that are recognized by multiple mice.

## 3 Materials and Methods

### 3.1 Mice and *in vivo* tumor treatment

The treatment model used here was previously described in detail (Baniel et al., 2020; Morris et al., 2016; Morris et al., 2018). In brief, B78-D14 (B78) tumor bearing mice were treated when tumors reached ∼ 100 mm^3^ with a combination of 12 Gy local radiotherapy (RT), followed 5 days later with 5 daily intratumoral (IT) injections of the hu14.18-IL2 immunocytokine (IC). Mice that were cured were rechallenged after 90 days with an additional injection of the B78 tumor. Mice that rejected the rechallenge were considered immune (**Figure 1A**). At indicated timepoints (**Figure 1A**), blood via mandibular bleed was collected into BD serum collection tubes and serum was harvested. For select animals a terminal bleed was obtained via cardiac puncture immediately following euthanasia to obtain larger volumes of serum from immune mice. Experiments were performed under an animal protocol approved by the Institutional Animal Care and Use Committee. A list of all serum samples from individual naïve and immune mice, used to generate the data presented in this report is included as **Supplemental Table 1**.

**Figure 1.**
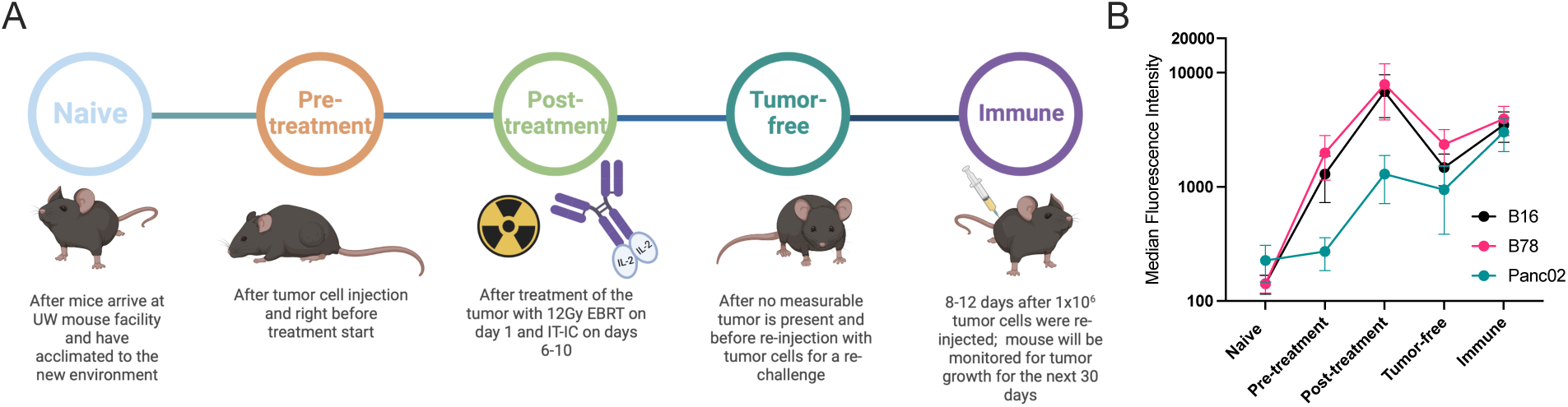
Mice develop antibodies against melanoma tumors throughout treatment. **A:** Timeline of blood serum collection. C57BL/6 purchased from vendors are allowed to acclimate 1-2 weeks prior to <u>Naïve</u> sample collection and B78 tumor implantation. After measurable tumors have established, <u>Pre-treatment</u> samples are collected prior to initiation of radio-immunotherapy [12 Gy external beam radiotherapy (EBRT) and intratumoral hu14.18-IL2 immunocytokine (IT-IC)]. Following completion of therapy, <u>Post-treatment</u> samples are collected. <u>Tumor-free</u> samples are collected from animals that have no palpable tumors ∼30 days following treatment initiation. ∼90 days post treatment initiation, these “cured” animals are rechallenged with tumor cells and <u>Immune</u> samples are collected the following week. Schematic created using BioRender. **B:** Flow cytometric analysis of serum antibody binding to tumor cells. Murine blood serum was incubated with murine tumor cells prior to staining with fluorescently tagged anti-mouse IgG antibodies and flow cytometric analysis. Median fluorescence intensity values corresponding to the timepoints described in A are shown. Serum samples were tested against B16 melanoma (black), B78 melanoma (pink) and Panc02 pancreatic adenocarcinoma (green) murine tumor cell lines. Error bars show standard error of the mean, n=3 mice for each datapoint.

### 3.2 Tumor cells

B78-D14 [“B78”, obtained from Ralph Reisfeld (Scripps Research Institute) in 2002] melanoma is a poorly immunogenic cell line derived from B78-H1 cells, which were originally derived from B16 melanoma (Becker et al., 1996; Haraguchi et al., 1994; Silagi, 1969). B78-D14 cells lack melanin, but were transfected with functional GD2/GD3 synthase to express the disialoganglioside GD2 (Becker et al., 1996; Haraguchi et al., 1994), which is overexpressed on the surface of many human tumors including melanoma (Nazha et al., 2020). B16-F10 melanoma was obtained from American Type Culture Collection (ATCC) in 2005. The murine pancreatic ductal adenocarcinoma cell line Panc02 was purchased from ATCC. Panc02, B78 and B16 cells were grown *in vitro* in RPMI-1640 (Mediatech) supplemented with 10% FBS, 2mMol L-glutamine, 100U/ml penicillin, and 100μg/ml streptomycin. Mycoplasma testing via PCR was routinely performed.

### 3.3 Flow cytometry

0.5×10^6^ cells of B16, Panc02 or B78 were used per tube and incubated with 1μl of serum for 45minutes. After incubation, cells were washed with 3ml flow buffer (PBS with 2% FBS) at 300xg and stained with goat anti-mouse IgG-APC (BioLegend, clone Poly4053, catalog # 405308) and rat anti-mouse IgM-PE (ThermoFisher, clone eB121, catalog # 12-5890-82) polyclonal antibodies. Cells were washed again at 300xg for 5min with 3ml flow buffer and resuspended in 50-100μl flow buffer. A drop of DAPI (BioLegend, catalog # 422801) was added to each tube before data was acquired on a ThermoFisher Attune flow cytometer. Data analysis was performed using the software FlowJo version 10.

### 3.4 High-density peptide array

#### 3.4.1 Design of mouse whole proteome peptide microarray

The mouse whole proteome peptide microarray was designed based on the protein set downloaded from UniProt in December of 2018 for C57BL/6 mice (The UniProt, 2017). The library was generated *in silico* for synthesis on high-density peptide microarrays (Nimble Therapeutics, Madison WI). The library consisted of overlapping 16-mers representing the entire mouse proteome tiled at every second amino acid for reviewed proteins and every 4 amino acids for most unreviewed proteins. All redundant (non-unique) peptides were only printed once but later computationally mapped back to all UniProt IDs containing this peptide. The individual peptides in the library were randomly assigned to positions on the microarray to minimize the impact of spatial biases.

#### 3.4.2 Peptide array sample binding

Mouse serum samples were diluted 1:100 in binding buffer (0.01M Tris-Cl, pH 7.4, 1% alkali-soluble casein, 0.05% Tween-20). Diluted sample aliquots were bound to arrays overnight for 16–20 hours at 4 °C. After binding, the arrays were washed 3x in wash buffer (1x TBS, 0.05% Tween-20), 10 minutes per wash. Sample binding was detected via goat-anti-mouse IgG Alexa Fluor 647 conjugated polyclonal antibody (Jackson ImmunoResearch, 115-605-071). The secondary antibody was diluted in secondary binding buffer (1x TBS, 1% alkali-soluble casein, 0.05% Tween-20) and incubated with arrays for 3 hours at room temperature, then washed 3x in wash buffer (10 minutes per wash) and 30 seconds in reagent-grade water. Then the array was washed 2x for 1 minute in 1x TBS and washed once for 30 seconds in reagent-grade water. Fluorescent signal of the secondary antibody was detected by scanning at 635 nm at 2μm resolution and 25% gain, using a micro-array scanner. Data were reported as arbitrary fluorescence units.

#### 3.4.3 Peptide array data processing

The datasets generated and analyzed for this study can be found on Zenodo under the following DOI: 10.5281/zenodo.7871566.

For each serum sample, the fluorescence intensity data from a single chip, for each unique peptide, was assayed and processed once; then results from identical peptides redundant to multiple proteins (i.e., were present in more than one protein represented) were restored to each protein. Raw fluorescence intensity signals from primary antibodies binding to peptides on the array, and secondary antibodies with a fluorescent tag binding to primary antibodies were reported. The amount of fluorescence signal is influenced by both the titer and affinity of primary antibodies binding to each peptide sequence.

#### 3.4.4 Data analysis workflow/pipeline of whole proteome data

Detailed bioinformatic/biostatistical data modeling, algorithms, analyses, and graphic presentation methodologies are beyond the scope of this manuscript focusing on the biology and immunology of what is detected using the sera of these immune mice. These issues, and their justification/rationale are presented in detail in a separate manuscript (McIlwain et al., 2023).

### 3.5 JPT peptide array

Samples were sent to JPT (JPT innovative Peptide Solutions, Berlin, Germany) and a custom designed PepStar Multiwell Peptide Microarray was performed following manufacturers protocol using a manufacturing process based on SPOT synthesis as described previously (Nahtman et al., 2007; Zerweck et al., 2016). Peptides were chosen based on different criteria from the high-density peptide array results, as described in the results section. We included 376 16-mer peptides with a range of signal from the high-density peptide array data and tested those on the same serum samples as well as additional serum samples from immune and naïve mice at a dilution of 1:100. Raw data obtained by JPT for these analyses were sent to us for further analysis and processing. Data were reported as arbitrary fluorescence units.

### 3.6 Peptide ELISA

For the peptide ELISA, 16 separate JPT BioTides^TM^ Biotinylated Peptides were purchased containing a TTDS-linker and biotinylation at the N-terminus. The peptides were generated using the same SPOT synthesis as the larger peptide array (Nahtman et al., 2007). Peptides were synthesized from C- to N-terminus ensuring that only full-length peptides will have a biotin at the N-terminus. Coating of streptavidin plates was performed per manufacturers instruction with a 250-fold dilution of lyophilized BioTide peptides. ELISA was performed according to JPTs peptide ELISA protocol with the adaptation to a 384 well plate instead of the standard 96 well plate to conserve on serum samples. Neutravidin coated 384 well plates by ThermoScientific (#15400) were used. Stop solution was added after a 30-minute TMB incubation. Plates were read at regular intervals during TMB substrate incubation (reads at 655 nm) and right after addition of stop solution (reads at 450 nm). Optical density values were used to analyze results.

### 3.7 Choosing of peptides for JPT and ELISA

Peptides for JPT analysis were chosen before the second dataset of whole proteome data using the high-density peptide array was generated and analyzed. 376 peptides were chosen based on different signal strength and reactivity to sample types. In more detail, peptides were chosen based on high signal (>500) in at least one immune sample and low to no signal (<20) in naïve samples. Some peptides were included because they shared, or partially shared, amino acid sequences. Others were chosen because they exhibited no antibody binding in any tested sample or because they had binding in every single sample. We also chose some peptides that exhibited low to medium signal.

For ELISA validation we chose a total of 16 peptides, 2 peptides without any reactivity in any tested sample, 5 peptides based on good correlation between JPT and whole-proteome results and 12 peptides (3 of which were also included in the JPT to whole-proteome correlation category) that showed significant binding in at least 3 immune samples in the moderate (or restrictive) category. We also confirmed that the expected binding sequence within each of these 16 peptides did not have the same, or very similar, sequence to those of any of the other peptides in this group of 16 peptides, to help ensure that each peptide would be identifying relatively distinct antibodies. We also chose 10 random peptides from all unique peptides from the array utilizing a random number generator.

### 3.8 Statistical analysis

#### 3.8.1 Peptide array processing

Data from 13 total unique serum samples were tested in the high-density microarray: 5 from naïve mice, 6 immune samples were obtained from mice following their RT+ IC induced cure from their initial B78 tumor, and then 8-12 days following their rechallenge with another injection of B78 tumor; and two samples (replicates) were obtained from separate mice after a 2^nd^ rechallenge injection of B78 tumor (**Figure 1A**). These 13 serum samples were assayed for antibody binding to 6,090,593 unique sequence probes mapped to a total of 8,459,970 unique probe IDs (due to redundancies in tiling across protein sequences and using a mixed tiling of either 2aa or 4aa across each protein), or a total of 53,640 individual proteins. Using spatially corrected processed data from Nimble Therapeutics, the data were log2 transformed, quantile normalized, and further processed using a sliding average mean window across the protein location of +/-8aa.

HERON (Hierarchical antibody binding Epitopes and pROteins from liNear peptides) (McIlwain et al., 2023) was developed and used to determine thresholds for calling antibody binding at the probe, epitope (consecutive probes), and protein level for each sample using meta-analyses methods to summarize binding across subjects in the post-rechallenge condition. Briefly, 1) a global p-value was calculated using a z-test for each probe signal using all sample and probe values, and 2) a differential p-value was calculated between the average of the naïve samples and each individual post-rechallenge (Tumor-free) sample. The global p-value and differential p-value for each post-rechallenge sample were then combined using the Wilkinson’s max meta p-value method (Wilkinson, 1951). After correcting for false discoveries using the Benjamini-Hochberg (BH) method (Benjamini & Hochberg, 1995), the individual probes for each post-rechallenge sample are considered bound by antibodies if their false discovery rates (FDR) are below a threshold. Epitope regions were identified by applying the skater algorithm (AssunÇão et al., 2006) to identify groups of antibody-bound probes (spatially and across subjects), and epitope meta p-values were calculated using the Wilkinson’s max method on the 2^nd^ highest probe p-value. Protein p-values were calculated using Wilkinson’s min (or Tippet’s) method (Tippett, 1931). After correcting the epitope and protein p-values using the BH algorithm, the epitope and protein sample calls were made using an FDR cutoff. To avoid prioritization of peptides that may be due to spurious noise, singleton probe and epitope calls without calls of neighboring probes or if the singleton call was not present in repeat immune samples it was removed. The number of samples that were bound by antibodies for each probe, epitope, and protein were tabulated as K of N statistics (K = # of samples with antibody binding; N = total # of samples).

#### 3.8.2 ICC score

Statistical analysis was conducted using R (v. 4.1.1; R Core Team 2021) and the packages ‘lme4’ (v. 1.1.27.1; (Bates et al., 2015) and ‘specr’ (v. 0.2.1; (Masur, 2020) for computing intraclass correlation coefficients (ICC). To analyze the agreement in the high-density whole-proteome, JPT, and ELISA instrument readings among selected peptides, log-transformed readings/intensity was modeled and compared using linear mixed-effects models, in which individual samples and the instruments were modeled, respectively, as random effects, while tumor stage and peptide, when applicable, were modeled as fixed effects. Intra-mouse correlation and intra-instrument correlation were accommodated via random intercepts. The ICC was computed from the instrument random effect to estimate their share of variance in the log-transformed readings. An ICC of 0-0.5 is considered poor reliability; 0.5-0.75 is considered moderate reliability, 0.75-0.9 are considered good reliability; 0.9-1 are considered excellent reliability.

#### 3.8.3 Linear regression/r^2^ score

Simple linear regression was performed using GraphPad Prism (Version 9.5.0, 2022) and r^2^ values were reported. These values were used to describe how predictive the high-density peptide data for the same sample is of the JPT peptide data. The closer the r2 value is to 1, the more predictive the high-density peptide array value is of the JPT value.

#### 3.8.4 Test of proportions

A test of proportions was used to compare the portion of positive reactivity between different peptide groups at a threshold of 2 or higher for OD readings. A p-value of 0.05 was considered statistically significant. The proportion of reactivity in the randomly selected peptides (1 of 200 with OD >=2) were found to be significantly less reactive than the HERON validation set (3+ FDR 0.05, 48 of 240 with OD>=2), respectively.

#### 3.8.5 Hypergeometric testing

The ELISA data replicates were first averaged together. A threshold of ≧2 O.D. (optical density) units was used to call positive antibody binding for each ELISA data point. For each peptide, the fraction of peptides with antibody binding was calculated for the immune samples in the original and validated ELISA set and for the naïve samples in the validated set. A peptide was considered validated if 25% or more of the respective samples were found to be positive.

To calculate the likelihood of getting n antibody-bound (positive) peptides out of a random sample of 14 draws, a hypergeometric distribution was used to calculate the probability of getting at least *x* positive hits out of 14 random draws, where the total pool of peptides tested by high-density array has *K* positive peptides^1^. As the true fraction of positive peptides (*K /* 6090593) within the pool of ∼6 million possible peptides to test from the high-density array is not known, we simulated the calculated p-value from the hypergeometric using different proposed fractions of the total positive peptides within the whole set of ∼6 million peptides tested by Nimble.

### 3.9 Summary of supplemental material

Supplemental Figure 1 showcases the raw flow cytometry histograms used to generate Figure 1B.

Supplemental Figure 2 showcases an example of repeated samples about a year apart while Figure 2 showcases an example of a repeat sample about a day apart.

**Figure 2:**
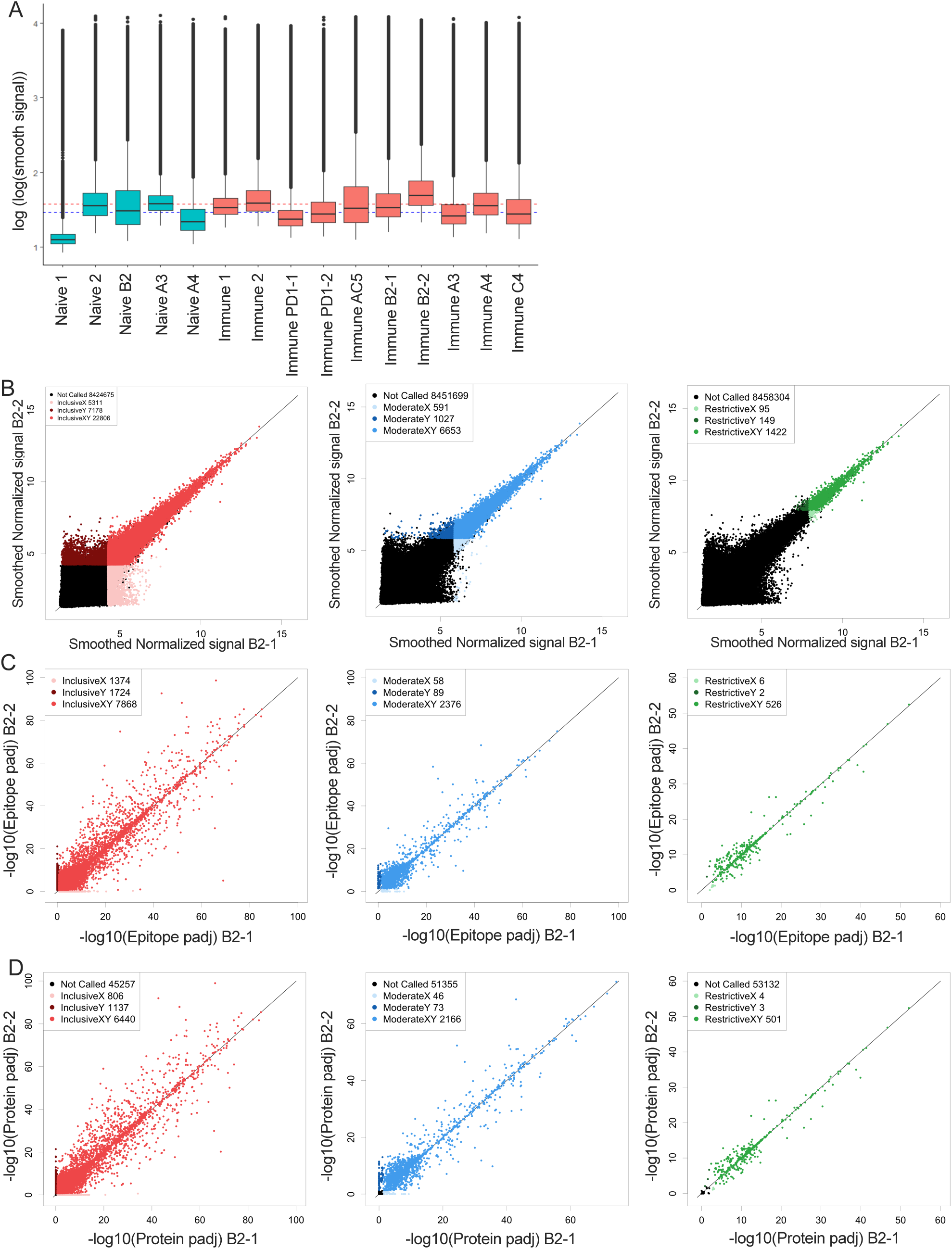
Overview of array data and reproducibility and reliability of probe, epitope, and protein calls. Serum samples from mice described in Figure 1A were run on the Nimble Therapeutics mouse whole proteome peptide microarray. **A:** Summary boxplot of signal intensity for sera from each of 6 mice, for which Immune sera were tested (designated: PD1, AC5, B2, A3, A4 and C4). For 3 of these mice, (B2, A3 and A4) naive sera results are also shown. Naive-1, and Naïve-2 are pools of sera from 6 separate naive mice (4 individual mouse samples per pool), and immune-1 and immune-2 are pools of Immune sera from 6 mice (4 individual mouse samples per pool). PD1-1 and PD1-2 are 2 replicate serum samples of cryopreserved Immune sera run independently, one year apart, on separate whole proteome microarray chips; B2-1 and B2-2 similarly are Immune sera from 2 replicate cryopreserved serum samples, run independently on separate whole proteome microarray chips within a day of each other. Data are presented as log-log transformed smoothed fluorescent intensities for all peptides in the array. Median values for all Immune (red) and Naïve samples (blue) shown are represented with dotted horizontal lines. **B:** Correlation of fluorescence intensity values from two separate whole-proteome microarray chips run one day apart (B2-1 and B2-2) on the same Immune-serum from one representative mouse. Each dot represents the log transformed processed raw array data for an individual called peptide. These peptides were then separated into 3 categories based on their signal strength: restrictive (highest signal, >10xSD above the mean), moderate (>6xSD above the mean), and inclusive (>3x SD above the mean) based on the statistical significance for each value above the mean signal strength for all peptides. Lighter colored dots represent peptide only called in the B2-1 assay, darker colored dots represent peptides only called in the B2-2 assay, for its respective category (Red: inclusive, Blue: moderate and green: restrictive). For each graph, the black dots are those peptides that were not called for that graph (at the indicated signal strength level) either in the B2-1 or the B2-2 assay. **C:** Scatter plots of epitope-level data based on the peptide data shown in Figure 2B, again segmented into restrictive, moderate, and inclusive rankings. Epitopes were identified based on overlapping consecutive recognized peptides and values plotted based on the -log10 p-values. Lighter colored dots represent peptides only called for sample B2-1, darker colored dots represent peptides only called for sample B2-2, for its respective category. **D:** Scatter plots of predicted protein-level data based on the peptide and epitope data shown in **Figure 2B&C**, segmented into restrictive, moderate, and inclusive rankings. Proteins were identified by combining epitope data and generating a protein p-value and values plotted based on the -log10 p-values. Lighter colored dots represent peptide only called in replicate sample B2-1, darker colored dots represent peptides only called in replicate sample B2-2, for its respective category. For **Figures 2 B-D** the numbers in the legend box within each figure indicate the number of dots in each category.

Supplemental Figure 3 shows epitope lengths across the different signal thresholds in addition to the restrictive category epitope length shown in Figure 3D.

**Figure 3:**
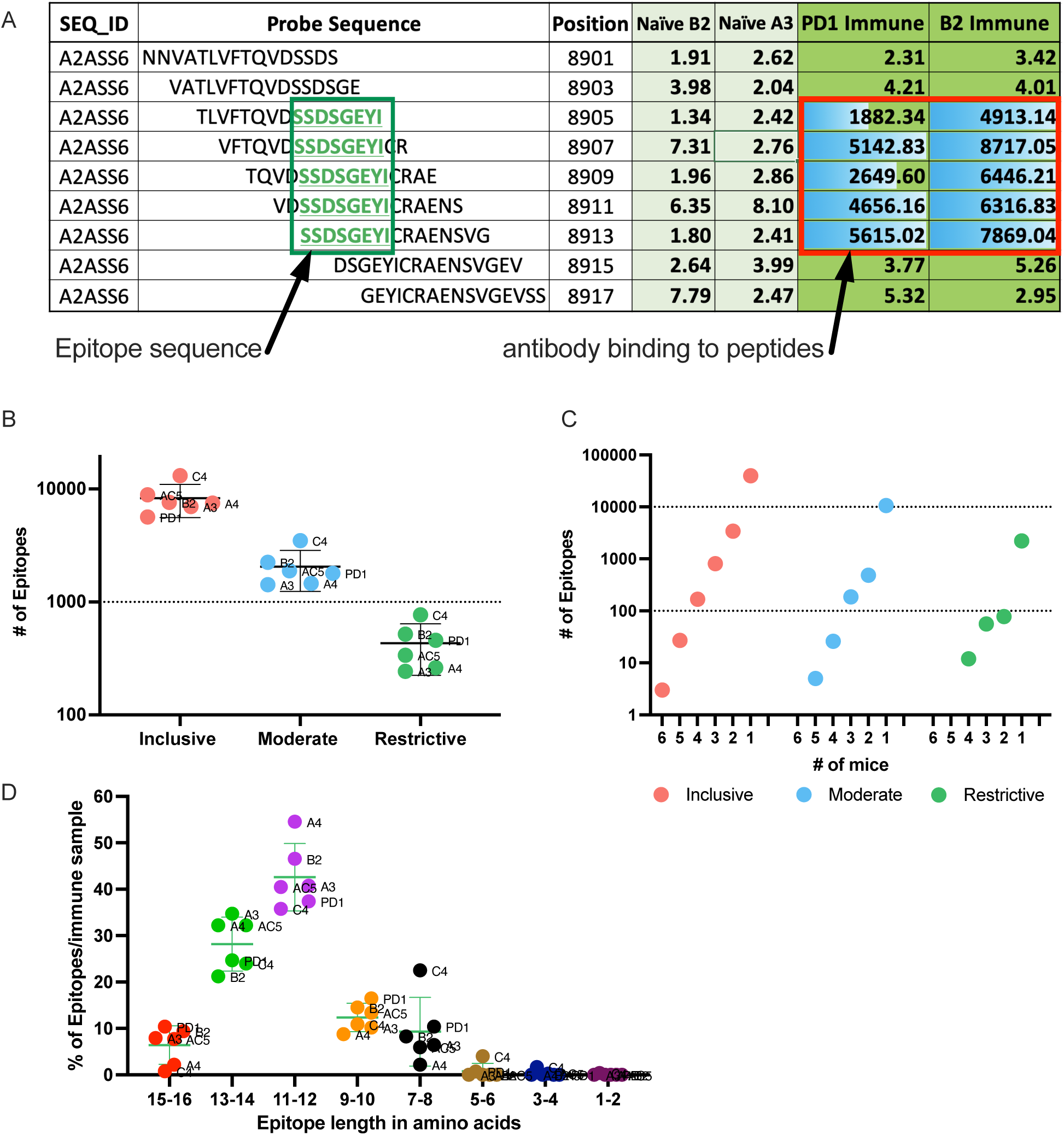
Number of epitopes identified and categorized from mouse whole proteome peptide microarray for all Immune samples. **A:** Example of raw data highlighting a predicted epitope, defined as a clustered and overlapping antibody binding region in the peptide microarray. A section of the titin protein is shown, with 9 stacked 16-mer peptides, each shifted by 2 aa positions, starting at aa position 8901-8917. Fluorescence intensity results are shown for each of these 9 16-mer peptides for separate serum samples from 2 naïve mice (naïve B2 and naïve A3) and 2 immune mice (PD1 and B2). Five of the consecutive 16-mers show strong binding by the 2 immune sera, while the other 4 16-mers show very weak binding by all 4 sera shown. The 5 well recognized 16mers each share the 8 sequential aa shown in the green box, indicating a recognized epitope **B**: Number of inclusive, moderate, and restrictive epitopes identified in the Immune samples with significantly higher antibody binding in Immune serum than in Naïve serum samples. Each dot represents the number of epitopes in that category, for each of the 6 separate mice tested. The individual mouse identifications are indicated next to each dot. **C**: Number of unique epitopes each recognized by any individual immune mouse, or co-recognized by 2, 3, 4, 5 or 6 Immune mice (of 6 total mice), segmented by cutoff category of inclusive, moderate, and restrictive of the epitopes. Within each category, the single dot plotted above the individual numbers plotted on the X axis indicate the number of epitopes recognized by exactly that number of mice. **D**: Categorization of epitopes by peptide length, based on the clustering as in Fig. 3A, using data from the restrictive category. Above each pair of numbers (i.e.: 1-2, 3-4, etc.) on the X axis are 6 colored dots each indicating the number of epitopes of that aa length recognized by each of the 6 mice tested.

Supplemental Figure 4 shows at what percentage of total positive peptides a significant p-value will be reached when utilizing different thresholds from our experimental values achieved from ELISA and validation ELISAs to extrapolate to the whole peptide array (∼6.5million datapoints vs 14 peptides tested via ELISA).

Supplemental Table 1 contains a list of all serum samples used in this manuscript.

Supplemental Table 2 provides additional information and calculations pertaining to Figure 2.

Supplemental Table 3 contains all peptides and samples tested in the JPT multiwell assay and their values that were shown or otherwise used for this manuscript.

Supplemental Data 1 contains all peptides and samples tested in the JPT multiwell assay and their corresponding high-density array values.

## 4 Results

### 4.1 Mice elicit a tumor-specific adaptive humoral response to RT+ IC treatment

B78 melanoma bearing mice treated with RT + IC + anti-CTLA-4 generated an antibody response to surface proteins on the B78 (or B16) tumor cells that was measurable at day 22 post tumor implantation (Baniel et al., 2020). To further investigate these antibody responses and to ensure that the RT+IC treatment alone (without added anti-CTLA-4) can elicit a similar antibody response, we collected serum at multiple times before, during and after successful RT+IC treatment of B78-bearing mice (**Figure 1A**). Serum was collected from mice at the following timepoints: before tumor cells were implanted (Naïve); once tumors reached treatment size but prior to treatment (Pre-treatment); within a week of mice completing the RT+IC regimen (Post-treatment); when the tumors were regressing but still present, weeks later after mice were deemed tumor-free and prior to a rechallenge (Tumor-free); and 8-12 days after subcutaneous rechallenge with injection of B78 cells, ∼90 days after treatment and >30 days after the mice were tumor free (Immune). At this point, a strong memory response was demonstrated based on the rejection of the rechallenged B78 tumors. These mice were monitored for an additional 5 weeks to ensure complete tumor clearance of the re-engrafted B78 tumor, proving that these mice were immune.

Using flow cytometry, we tested the serum from each of these timepoints for IgG antibody binding to B78 cells. We observed the presence of endogenous anti-tumor antibodies against B78 in tumor-bearing mice starting at the pre-treatment timepoint, with increasing levels of antibody detected by flow cytometry at all subsequent timepoints (**Figure 1B**). To determine the specificity of these anti-tumor antibodies, we also incubated these serum samples with B16 melanoma cells, the parental line to B78 that is GD2 negative as well as a separate syngeneic pancreatic adenocarcinoma cell line, Panc02. Serum antibodies showed recognition of B16 to a very similar degree as to B78 and a lower recognition of Panc02 (**Figure 1B**). Recognition of Panc02 cells by these serum samples might reflect some shared surface antigens between B78 and Panc02 cancer cell lines. The data presented in Figure 1 are the summed flow cytometry results for 3 of the 6 mice studied subsequently in the high-density peptide array, described below; individual mice showed slight variations in the strength of the responses to these 3 tumor lines at different timepoints (**Supplemental Figure 1**).

### 4.2 Whole proteome peptide array results are reliable and repeatable at high signal levels

To investigate what these antibodies are recognizing on the tumor cells, we used a whole proteome peptide array to profile antibody recognition comparing serum from the naïve vs. the immune timepoints (as shown in **Figure 1A**). First, we plotted the signal for each of the 6 mice against all 8.46 x10^6^ individual peptides, referred to as probes, as a boxplot for each individual sample (**Figure 2A**). The overall appearance of immune and naïve samples is very similar, with most probes giving a signal near the baseline, and a small fraction of probes giving signals 100-1000 fold higher than baseline. Even so, more of the probes (detailed numerically in the next paragraph) have even stronger signals in the immune sera, such that the mean of all immune samples is greater than the mean of all naïve samples (**Figure 2A**). Our hypothesis was that specific peptides would show significantly higher binding in immune samples compared to naïve samples. Overall, since we are measuring antibody responses to all native peptides within the mouse proteome, we did expect to see antibody binding to some of these peptides in naïve as well as immune samples as previously seen by Hulett *et al* in 2018 (Hulett et al., 2018).

Prior to identifying high binding peptides recognized by individual or multiple mice, we evaluated how reproducible the signal strength is using this high-density peptide array system. Serum samples from an individual immune mouse (mouse B2), taken after rejection of rechallenge (the immune timepoint in **Figure 1A**) was divided and separate aliquots were analyzed in the same array assay, on independent “chips”, each quantifying the binding signal against all 8.46×10^6^ 16-mer peptides. The paired values for each of these peptides, in the 2 parallel samples are plotted on the X and Y axes in **Figure 2B**. We first looked at all peptides with significantly higher binding than the mean overall signal, defined as a signal that is larger than three standard deviations (SDs) above the mean (inclusive) (left panel of **Figure 2B, red**). Although difficult to appreciate due to the number of data points overlying each other, there are 8,424,675 black data points (corresponding to both of the values for that peptide being ≤3SD from the mean value), and only 35,295 non-black data points in the Inclusive group, indicating that at least one of the data points was >3SD (**Figure 2B, red)**. A similar analysis was performed for 2 separate aliquots of immune serum from the same blood sample, but from a separate immune mouse (mouse PD1), that was performed on 2 separate identical chips against all peptides, but the analyses were run on separate days, ∼ one year apart (**Suppl**. **Figure 2A**). As for **Figure 2B**, there are 8,423,302 black data points, and only 36668 non-black data points (Suppl. **Figure 2A)**. However, when looking at all probes that fit these criteria and plotting replicate sample results against each other (**Figure 2B** and Suppl. **Figure 2A**), we noted that at 3SDs, we found a number of probes where one of the values was >3SD from the mean but where the replicate sample gave a result that was ≤3SD from the mean. In **Figure 2B**, these are the probes shown off the diagonal in light-red or dark-red, while all the probes shown along the diagonal in red correspond to *<u>those where both values</u>* from the two separate “runs” of the same serum sample gave concordant values >3SD from the mean. In **Figure 2B**, the number of probes in red, (i.e., seen by both replicates, and designated in the legend box as “InclusiveXY”), corresponds to 65% of the non-black probes with 35% of the non-black probes comprised of the lighter and darker red probes (**Figure 2B**). Because one of the values for these lighter and darker red probes was not >3SD from the mean, these values were not consistent or reproducible, therefore less reliable to call as antibody binding hits.

To enhance reliability and reproducibility of results, we increased the signal strength criteria to ≥6SDs above the mean (middle panel of **Figure 2B**, moderate, in blue) which included the top ∼0.1% of peptides compared to the top 0.4% of peptides at 3SDs in the inclusive category. This moderate category showed a larger concordance of recognition between the two replicates, with 80% of the non-black probes being identified at this level by both samples, shown in blue (6653 probes); only 20% of the probes showed discordant signals (one sample >6SD with the paired sample being ≤6SD) in the lighter (591 probes) and darker blue (1027 probes). The best reproducibility between the 2 paired samples on a peptide level was achieved with the restrictive category which was set at 10SDs (right panel of **Figure 2B**, restrictive, green), which includes only 0.02% of all peptides. At 10SDs, over 85% of probes that are not black (at least one of the 2 values >10SD) were in the green category, with both probes >10SD (1422 probes) while only 15% of probes showed discordant signals (one sample >10SD with the paired sample being ≤10SD), in the lighter (95 probes) or darker (149 probes) green. The specific numbers of probes in each category, for each of the paired serum samples for figures 2B and Suppl Fig. 2A are provided in Supplemental Table 2.

We developed the HERON algorithm to identify consecutive overlapping, reproducible probes with high signal, and categorized the shared aa sequences represented by those highly recognized probes as epitopes based on specified thresholds (**Figure 3A**). The mean signal of an epitope was calculated based on the mean signal of all peptides that comprise the epitope. We again used the categories of inclusive, moderate, and restrictive (based on the single probe calls, but now based on standard deviations as well as false discovery rates (FDRs) to assure that all epitopes with significant signal were counted. Reliability and reproducibility for each of these categories increased significantly by looking at epitopes rather than probes alone and helped eliminate many of the non-reproducible binding events seen only on X or Y axes, but not both (**Figure 2C** and **Supplemental Figure 2B, Supplemental Table 2**). From samples from the same run, the percent of epitopes in the inclusive group of epitopes co-recognized by both samples improved from 65% (for peptides in **Figure 2B, Supplemental Table 2**) to 72% (for epitopes in **Figure 2C, Supplemental Table 2**); in the moderate category from 80% to 94% and in the inclusive category from 85% to 98.5% (**Figure 2C, Supplemental Table 2**). When looking at data from separate runs of the same sample, it improved from 37% on the inclusive probe level to 43% on the epitope level. For the moderate category, it increased from 49% to 63%, and in the restrictive category, it increased from 53% to 73% (**Supplemental Figure 2B, Supplemental Table 2**).

We further assessed reactivity of the sera to which protein these peptides and epitopes could correspond to based on specific criteria. Proteins recognized by immune sera were defined as proteins containing epitopes recognized by the immune sera, using the same criteria for defining inclusive, moderate and restrictive categories of proteins based on the epitope signal for the strongest epitope signal in that protein being either >3, 6 or 10 x SD significantly greater than the mean at FDR adjusted levels of 0.2, 0.05, and 0.01 respectively, clustering across probe hits using skater (AssunÇão et al., 2006) to find epitopes, and filtering regions that are likely spurious signals reflecting individual probes with a positive signal, but without a signal for the flanking peptides, and rescoring using the Wilkinson’s 2^nd^ highest max and the Wilkinson’s min/Tippets (Dewey, 2022; Tippett, 1931; Wilkinson, 1951) for the epitope and protein p-values respectively and using Benjamini Hochberg to calculate adjusted p-values on the probe, epitope, and protein level before making calls at the corresponding FDR levels (Benjamini & Hochberg, 1995)(as shown in **Figure 4A**). At the protein level, we again were able to see an increase in reliability of called proteins based on signal strength (**Figure 2D**). The stronger a protein was recognized (based on signal strength within each epitope in the protein), the higher the percentage of co-recognition by the 2 replicate serum assays (plotted on the X and Y axes) were found: the inclusive category showing 78%, moderate category showing 95% and restrictive category showing 99% of proteins co-recognized by both replicate samples (**Figure 2D, Supplemental Table 2)**. When comparing data from the same sample from different runs, the percent of proteins co-recognized by both replicate samples also increased in comparison to probes and epitopes, with 50% in inclusive, 66% in moderate and 75% in restrictive (**Supplemental Figure 2C, Supplemental Table 2**). Overall, we see increasing levels of reproducibility (namely co-recognition of the same peptides, epitopes or proteins by the 2 replicate samples evaluated on separate chips on the same day, or on different days), when going from the inclusive to the moderate to the exclusive category. Furthermore, we see increasing levels of reproducibility within each of these 3 categories, when going from peptide to epitope to protein.

**Figure 4:**
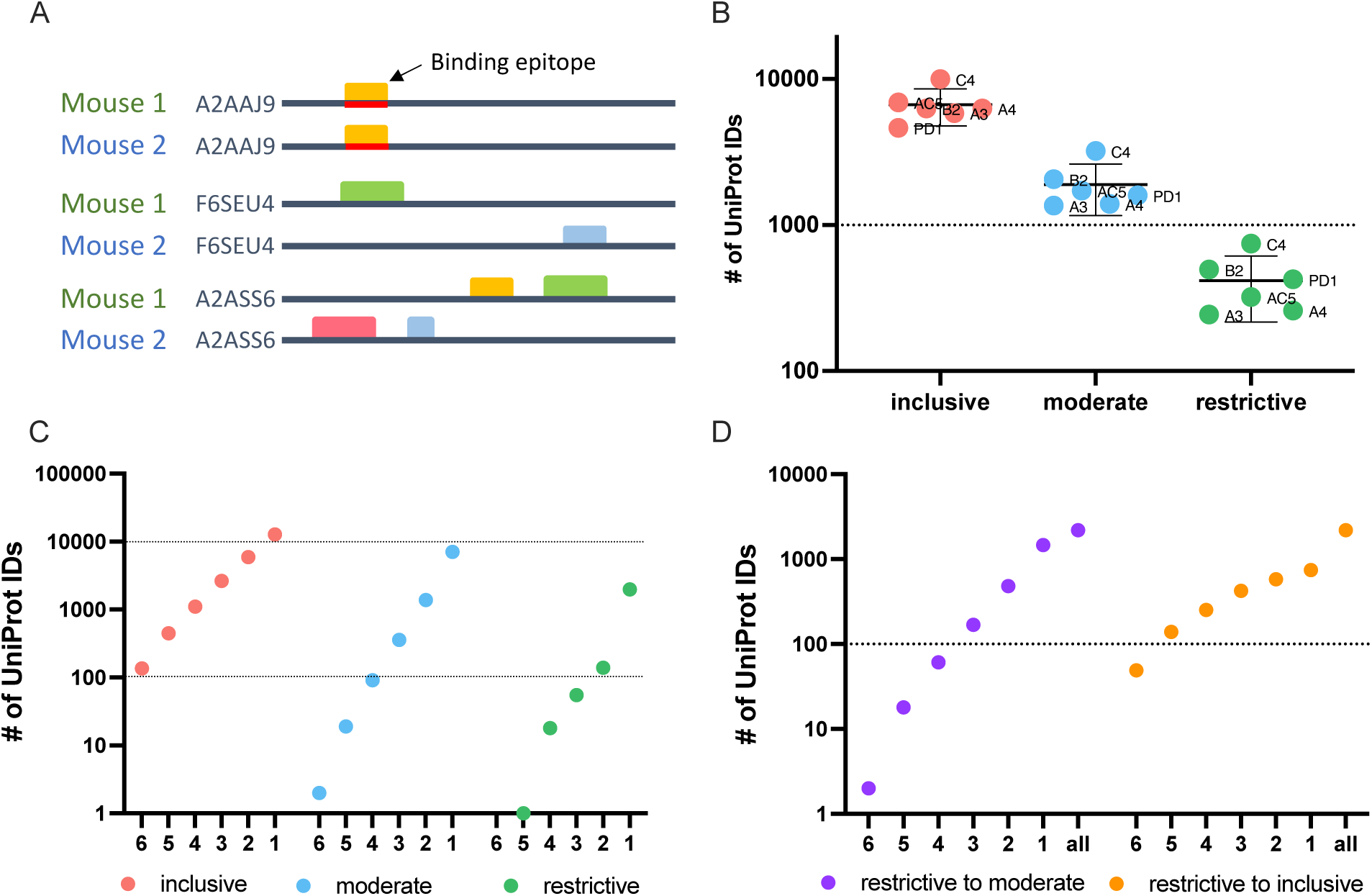
Protein level analysis of Nimble Therapeutics mouse whole proteome peptide microarray data and identified epitopes. **A**: Schematic showing examples of conditions that can lead to identification of one protein recognized by sera from 2 separate mice, including conditions where the same epitope within the protein is not recognized by antibodies from both mice, even though the protein is recognized by sera from both mice. **B**: Number of unique UniProt IDs recognized per Immune sample, segmented by the inclusive, moderate, and restrictive categorization of the called protein. Mouse IDs are labeled on each dot to demonstrate similar overall distribution within each category. **C**: Number of inclusive, moderate, and restrictive unique proteins recognized by at least 1, 2, 3, 4, 5 or 6 of the 6 mice tested. **D**: Number of proteins identified by the given number of mice on the X axis, where at least one mouse recognized the protein within the restrictive category and the other mice identified that same protein at least in the moderate category (purple dots) or at least in the inclusive category (orange dots) in the other samples.

### 4.3 Some epitopes are identified by multiple mice

Using consecutive peptides that show high fluorescence signals in immune sera but not in naïve sera enabled us to identify binding epitopes as well as which part of the peptides contained the binding sequence. **Figure 3A** shows at the top an exemplary 16 aa sequence of the protein Titin(UniProt ID A2ASS6), ranging from aa position 8901 to position 8917. Under it are 8 more consecutive 16-mer peptides, each shifted 2 positions to the right from the one above it, thus overlapping with it by 14 amino acids. For this section of Titin, no binding was observed to any of these 9 peptides by the 2 naïve sera tested (A3 & B2). However, strong binding was seen, reflected in high signal numbers, by sera from 2 immune mice, PD1 and B2. The center 5 peptides all show strong binding by both of the immune sera, indicating that the shared 8 aa sequence of these 5 peptides, SSDSGEYI, reflects the antibody binding sequence. The shared 8 aa sequence, recognized in these 5 overlapping peptides, is referred to as an epitope. Overall, using data from the high-density peptide array, we were able to identify an average of 6400 epitopes in the inclusive category, 2200 epitopes in the moderate category and just under 500 epitopes in the restrictive category by the immune serum samples from each of the 6 immune mice studied (**Figure 3B**). Of the identified epitopes, many were recognized by only one mouse, while some epitopes were recognized by sera from 2 or more mice, with one epitope being recognized by sera from all 6 mice in the inclusive category (**Figure 3C**). However, with increasing signal strength requirements, that same epitope was seen by fewer than 6 mice when using the moderate or restrictive category. In the highest binding (restrictive) category, twelve epitopes are each recognized by sera from 4 mice (**Figure 3C**), while 2450 of 2644 epitopes are recognized by only a single mouse (different epitopes for different mice). For the moderate category, 11491 of 12327 epitopes are recognized by only a single mouse. In the inclusive category, 46493 of 51664 total epitopes are recognized by only a single mouse. These findings are in line with previous studies looking at protein arrays where the abundant and heterogeneous nature of plasma and serum autoantibodies, regardless of disease status, was discussed (Ayoglu et al., 2013; Nagele et al., 2013).

Previous reports stated that an average length for a linear B cell epitope is around 5 to 12 amino acids (Buus et al., 2012; Engmark et al., 2016; Kringelum et al., 2013). Consistent with this, over 90% of epitopes within the restrictive category are between 7 and 16 amino acids long (**Figure 3D).** Note that a very small fraction of epitopes is identified with a length of 1-2 aa. These small epitopes may be an artifact of the computer algorithm, and most likely suggest that at least two separate antibodies in an individual mouse’s serum are binding to overlapping epitopes in this 1-2 aa region, such that we are actually measuring the overlap of the 2 longer epitopes. However, with the data that we have, it is impossible to determine the start and end for each individual overlapping epitope within the region. In general, we found that epitope length varies slightly across binding strength categories, an increase in shorter epitopes is visible in the lower categories (**Supplemental Figure 3A & B**).

### 4.4 A greater fraction of proteins than epitopes are bound by sera from multiple mice

A greater fraction of recognized proteins was bound by sera from multiple mice than were found when evaluating epitopes. This difference in proteins vs. epitopes recognized by multiple mice reflects the different requirements for the determination of recognition of a protein vs. an epitope. For an epitope to be recognized by sera from 2 separate mice, the 2 serum samples need to recognize the same epitope. In contrast, for a protein to be recognized by sera from 2 separate mice, each of the 2 serum samples need to recognize that protein, but not necessarily at the same place on the protein; in other words, if the 2 serum samples recognize distinct epitopes, even at opposite ends of the protein, then these 2 serum samples still recognize that individual protein, as shown schematically in **Figure 4A**. For each of the 6 mice tested, an average of 5089 recognized proteins were in the inclusive category as compared to 1963 in the moderate and 447 in the restrictive category (**Figure 4B**). However, using sera from multiple mice, 4323 proteins were recognized by at least 3 mice within the inclusive category, but only 136 proteins were found to be recognized by sera from all 6 mice. In the restrictive category, 74 proteins were recognized by sera from at least 3 of 6 mice, and 469 proteins from the moderate category were recognized by at least 3 of 6 mice (**Figure 4C**).

To broaden the criteria for recognition by sera from multiple mice, we focused on all proteins that were detected in the restrictive category by at least one mouse (2188 total unique UniProt IDs) and looked at these UniProt IDs to see if they were detected by sera from any of the other 5 mice using the restrictive to moderate (purple) criteria (**Figure 4D**) to see how many of these mice would recognize these same proteins when the signal strength requirement was loosened. We were able to detect 2 proteins that were now recognized by all 6 mice in this restrictive to moderate category. Overall, 33% of proteins seen by at least one mouse using the restrictive category were seen by 2 or more mice, and 11.4% were seen by 3 or more mice. A similar analysis is also shown for proteins recognized by at least one mouse in the restrictive category, and by other mice using the inclusive (orange) criteria (**Figure 4D**). This showed 66% of proteins seen strongly by at least one mouse are recognized by two or more immune mice, while 40% are recognized by 3 or more mice. These analyses indicate that there are several proteins recognized by more than one mouse, while the strength of the recognition signal of the peptide array system can vary from mouse to mouse.

### 4.5 Separate peptide ELISA techniques validate whole proteome peptide array data

After we established HERON, the method used above for the detailed analyses of peptide array data of the proteome recognized by immune sera from mice (and detailed further in a separate companion bioinformatic manuscript (McIlwain et al., 2023), we wanted to validate our findings with a separate, independent, antibody detection system for 16-mer aa probes that uses a different technology. For this we used a JPT multi-well peptide array to test 189 16-mer peptides per slide, allowing for testing of a larger number of serum samples. We chose peptides to use in this JPT system based on the results obtained using HERON analysis applied to data from the analyses of the entire proteome, summarized above. We chose to include a small number of peptides that showed no binding in any (naïve and immune) serum samples and chose a larger number of peptides that showed significant level of binding by one or more of the immune serum samples using the data from the proteome analyses from the 6 immune mice tested. The full panel of peptides selected, and the level of their reactivity with naïve and immune serum samples are presented in **Supplemental Table 3**. We used some of the same serum samples that we previously tested on the whole proteome high-density array (**Figure 5A**) to test these 189 peptides on the JPT array (**Figure 5B**). We show the mean reactivity for these same peptides and these same sera using the whole proteome data and the JPT system data **(Figure 5A & 5B).** Both naïve samples show no binding in either the high signal peptide or no signal peptide groups on the whole proteome peptide array as well as the JPT multi-well peptide array (**Figures 5A, 5B**). Immune serum samples showed very similar trends, with higher mean signals seen for the high signal peptides than for the no signal peptides. The A3 and A4 immune samples have a low mean signal for the high signal peptides in the whole proteome array as well as JPT. Overall, these results show that Nimble peptide array data can be qualitatively reproduced using an independent JPT multi-well peptide array. Note that the peptides in the Nimble system are biotinylated at the opposite end from that for the JPT peptides, and thereby fixed to the plate at opposite ends; this makes the peptide available to the sera in reverse orientation, thereby partially accounting for non-identical recognition patterns for the same peptides in these two systems.

**Figure 5:**
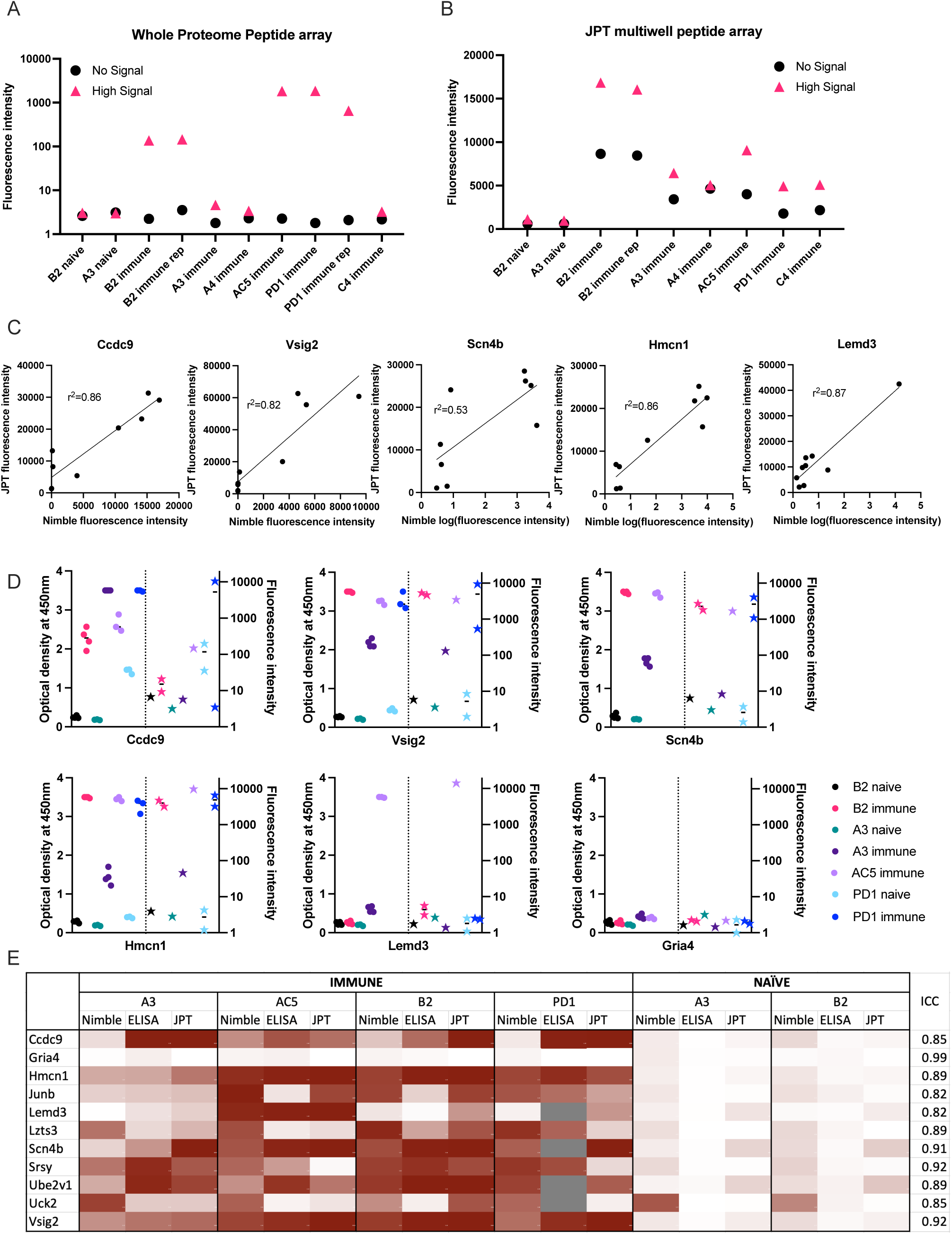
Comparisons of data from Nimble and JPT systems, for the same peptides and serum samples. **A & B:** Comparison of results using the same 10 serum samples tested in both JPT and whole proteome (Nimble) systems, for 11 peptides selected from the whole proteome data to show no significant signal with any serum samples (naïve or immune) vs. 272 peptides showing a high signal with at least one immune serum sample (A&B). **A:** Median fluorescence intensity values (from the whole proteome system) for peptides with a high signal (>1000 fluorescence units, >10SD over the mean) in at least 1 immune serum sample that were also tested on the JPT peptide array (pink triangle, high signal, 272 peptides) and on 12 peptides with a signal below 10 fluorescence units in all samples (identified based on the immune and naïve samples from the first whole proteome chipset) (black circle, no signal, 11 peptides) are displayed for 10 serum samples tested in the whole proteome system. **B:** Median values for the same peptides as shown in A are shown for the same samples (minus the repeat PD1 sample which was only run once on JPT) run on JPT multi-well peptide array. **C:** JPT (Y-axis) vs. Nimble (whole proteome, X-axis) fluorescence signal data-comparison plots for 5 exemplary peptides. **D:** Whole proteome data to ELISA comparison plots for 6 representative peptides (5 of which are the same as displayed in **C**) on 7 separate serum samples. For each peptide shown, the left Y axis shows ELISA data as optical density readings, and the right Y axis shows original whole proteome peptide array fluorescence intensity data for the same serum samples on the same peptide. The 7 individual serum samples are displayed in each graph with the same color, ELISA data are shown as circles, whole proteome data are shown as stars. The vertical dotted line separates ELISA from Nimble data. Multiple datapoints [dots (ELISA) or stars (Nimble, whole proteome) for one sample] show replicates. **E**: Heat map of 11 peptides from 11 different proteins with results from 4 immune serum samples and 2 naïve samples across 3 different peptide binding assays. Results from Nimble whole proteome peptide array as well as JPT peptide array and peptide ELISA were performed on the same serum samples and peptides; results are visualized via heatmap. Eight of the peptides shown were selected based on significant binding by at least 50% of immune serum samples in the whole proteome system. ICC: Intraclass correlation coefficient, is the reliability measure of the instrument for that specific peptide accounting for time of treatment and mouse. ICC scores of 0-0.5 show poor reliability,0.5 - 0.75: Moderate reliability, 0.75 −0.9: Good reliability and 0.9-1 excellent reliability.

The assessment of responses to some of the individual single peptides tested in both systems, demonstrates a qualitative relationship between the magnitude of responses by individual immune mouse serum samples, when tested on the same peptide in the Nimble and JPT systems (**Figure 5C**). We examined the same peptides recognized by the same serum samples as shown in **Figure 5C** utilizing a separate peptide ELISA system (**Figure 5D).** The ELISA data showed these same serum samples show a qualitatively similar pattern to that seen using the Nimble data for these same peptides. A summary of Nimble to JPT to ELISA comparison for 11 peptides using 4 immune and 2 naïve samples is shown in **Figure 5E**. The overall intraclass correlation coefficient (ICC) for instrument (Nimble, JPT, ELISA), considering peptide, tumor stage, and intra-mouse correlation, was 0.86. At the peptide-level, accounting for naïve vs. immune and intra-mouse correlation, these comparisons show 4 peptides with excellent ICCs (> 0.90: Gria4, Scn4b, Srsy and Vsig2) and 7 with good ICCs (0.75-0.90). None of the tested peptides received a moderate (0.5-0.75) or poor (< 0.5) ICC, thereby demonstrating that these 3 ways of measuring antibody responses are not important sources of variation in the measurement of antibodies to these peptides. Overall, these three assay systems showed similar patterns of response for the peptides we chose to evaluate.

### 4.6 Single peptides follow a similar trend in reactivity as seen with surface staining via flow cytometry

We chose 3 peptides to test all timepoints of serum collection shown in **Figure 1** using samples from 2 mice. These peptides were chosen based on high signals for these 3 peptides using most immune samples tested, as well as low signals with most naïve samples tested. **Figure 6A** shows how the level of antibody from mouse B2 towards the specific peptide increases with each subsequent serum sample (as detected by peptide ELISA) until reaching a plateau and then remains at that peak level while **Figure 6B** shows overall lower levels of antibody (as detected by peptide ELISA) for mouse A3 towards the selected peptides. Furthermore, we were able to observe stable antibody concentrations from post-treatment to tumor-free timepoints followed by an increase in antibody in our immune timepoints. In contrast to the flow data reactivity to B16 cells (**Figure 1B**), we were not able to see that the post-treatment timepoint exhibited the highest antibody binding for these three specific peptides.

**Figure 6:**
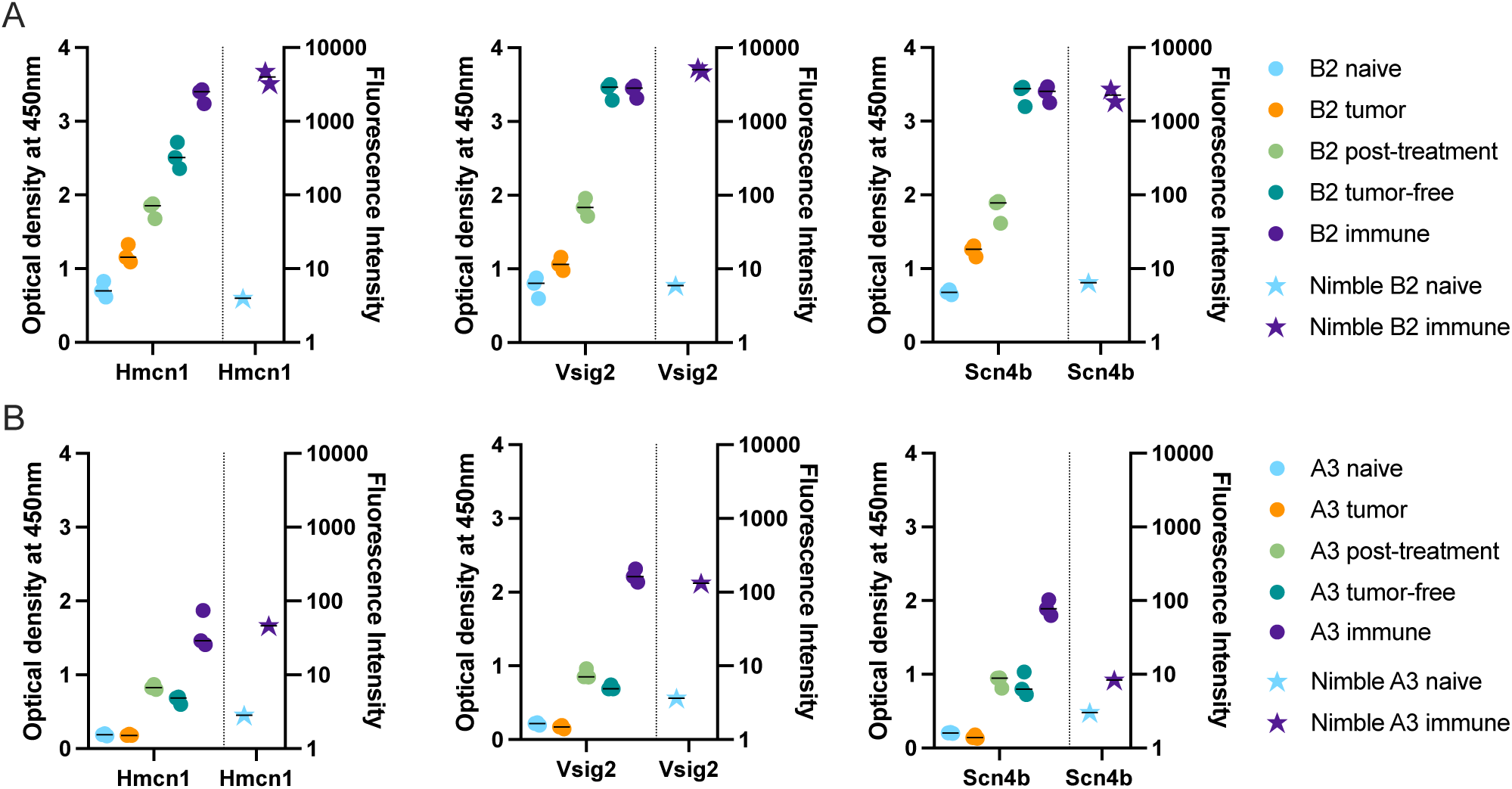
Time-course analysis and validation of Nimble peptide array results via peptide ELISA. **A&B**: Peptide ELISA of three exemplary peptides (16-mer peptides belonging to Hmcn1, Vsig2 and Scn4b) on all serum collection timepoints shown in Figure 1 on the indicated separate serum samples from 2 immune mice [B2, top row (**A**) and A3, bottom row (**B**)] are shown as optical density on the left Y axis. Right Y axis displays the corresponding fluorescence intensity from the Nimble Peptide array system for the indicated naïve and immune timepoints. Three separate replicate data points are shown for each serum specimen for each peptide in the ELISA (left Y axis), and 2 replicate data points are shown (at times these overlap) for each serum sample on each peptide for the immune Nimble (whole proteome) data (right Y-axis).

### 4.7 Validation cohort shows binding to most of the 14 peptides selected for binding in immune samples

Lastly, we hypothesized that peptides recognized by immune (but not naïve) sera by 50% or more of our initial cohort of 6 mice would also be seen by sera from additional similarly treated immune mice (as in **Figure 1A**) that were not ever previously tested on any peptide array or ELISA. To test this hypothesis, we selected the following peptides for ELISA testing: 1) 12 well-recognized peptides with binding by sera from at least 3 of 6 immune mice in the moderate category (**Figure 2B**) as tested in the whole proteome peptide array, 2) selected 2 peptides with significant binding in one or two of the original mice, and 3) 2 peptides without any significant binding in any serum samples from the Nimble system. Of the 6.09 x10^6^ unique peptides tested in the Nimble system for these 6 immune mice, only 316 peptides (0.005%) showed recognition at the moderate level for at least 3 of 6 immune mice. **Figure 7A** shows, by heat map, the original whole proteome data for these 14 recognized and 2 non-recognized peptides for 5 of the original 6 mice. We then tested these same 16 peptides via ELISA on the same naïve and immune samples as we had run on the whole-proteome peptide array (in **Figure 7A**) and obtained qualitatively comparable results (**Figure 7B**). We were able to test the same 5 mice but didn’t have enough serum left for all peptides with mice A4 or PD1. In this ELISA 12 of the 14 previously selected peptides show significant recognition by at least 1 immune mouse, and 8 of the 14 peptides are recognized by at least 2 of these 5 mice. We then proceeded to run these same 14 reactive peptides (and 2 non-recognized peptides) on new naïve and immune samples (previously untested by array or ELISA) collected from mice who had the same B78 tumor and received the same RT + IC therapy as our initially treated mice. ELISA results of the 20 new immune and 14 new matched naïve serum samples are shown in **Figure 7C**. Overall, we were able to show that ∼13 of the 20 new immune mice (65%) have antibodies against at least 1 of the 14 reactive peptides with 10 mice showing reactivity to multiple peptides. All new samples exhibited no antibody binding against the 16-mer peptides from Gria4 or P53, just like none of the original samples did in **Figures 7A&B**. However, 6 peptides showed binding to at least 1 naïve sample with 4 of the 14 naïve samples showing some antibody reactivity against at least 1 peptide (**Figure 7C**).

**Figure 7:**
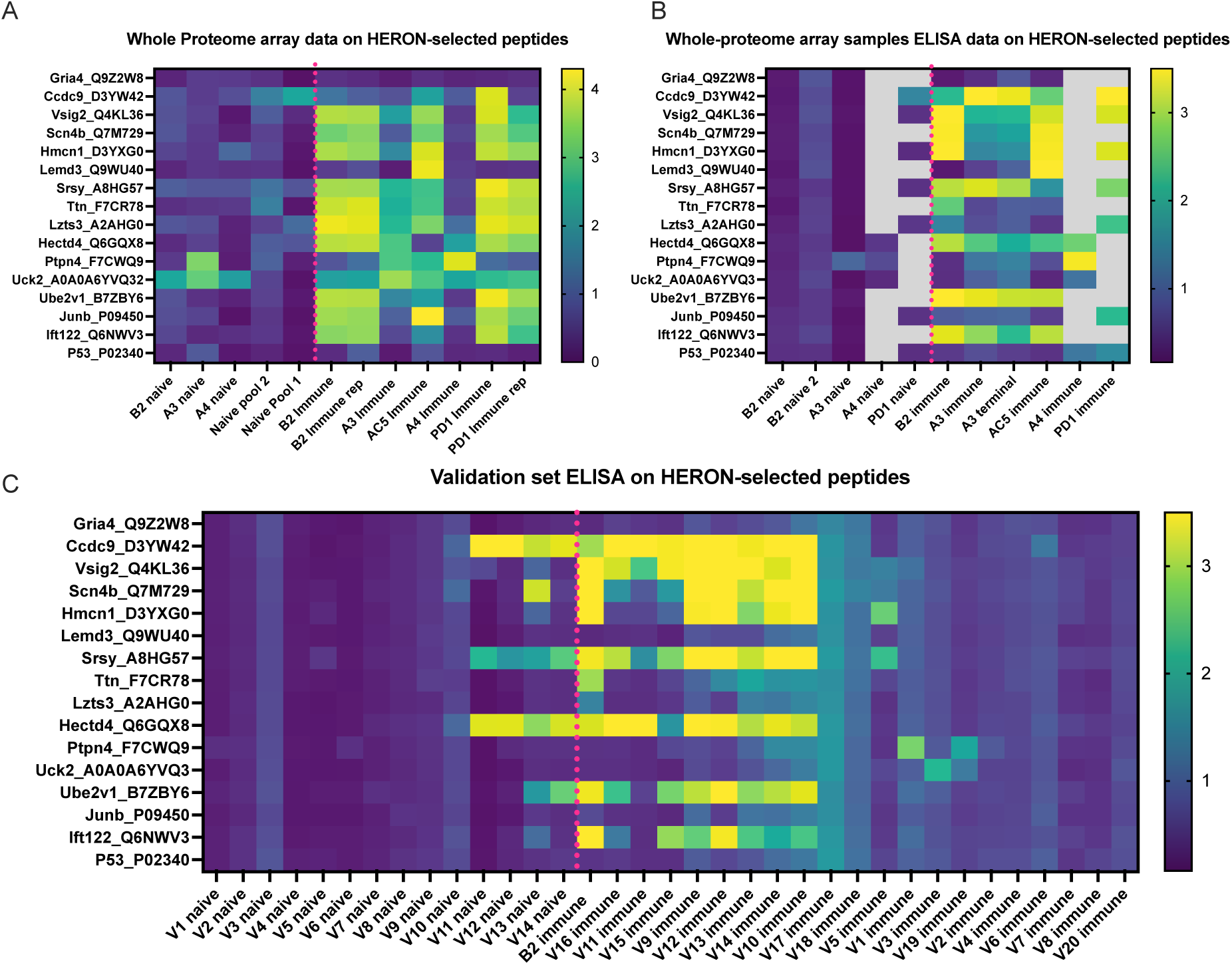
Peptides identified by the whole proteome array are also seen by ELISA testing for the same 6 immune mice, and for a separate validation set of 20 separate immune mice. **A:** Heatmap of 16 chosen peptides from whole proteome peptide array displaying whole proteome peptide array sample results for 12 serum samples including 2 replicate samples (B2 immune and PD1 immune). Dotted pink line separates naïve serum samples on left from immune serum samples on right. Twelve of these 16 Peptides were chosen, based on whole proteome data, demonstrating significant binding by serum samples from at least 3 of the 6 immune samples. Two of these peptides were chosen because of selective reactivity in only one or two of the original samples (Lemd3 and Ccdc9). Two of these 16 peptides shown were selected because they exhibited no binding by any of the immune or naïve serum samples tested in the whole proteome system (Gria4 & P53, at the top and bottom of the list shown). Data shown are log10 of the fluorescence units of the peptide array signal. **B:** Heatmap of ELISA results using the same peptides and serum samples as in Figure 7A. Grey areas indicate peptides not tested for the 4 indicated serum samples. Data shown are Optical Density (O.D.) values read at 450 nm length on a scale from 0 to 3.5. **C:** ELISA data for the same peptides as in A & B but using immune mouse serum samples never tested before from 20 separate mice that have received the same treatment to cure their B78 cancer (together with matched naïve serum samples for 14 of these 20 immune mice). Also included here is a repeat immune serum sample from one of the 6 immune mice used in the original whole proteome samples as an internal control (B2 immune, also shown in whole proteome data in Figure 7A, and ELISA data for original whole proteome samples in Figure 7B). Data shown are optical density values read at 450nm length on a scale from 0 to 3.5.

To contrast the large amount of antibody binding observed within this targeted selection of peptides we developed and utilized HERON to choose peptides recognized by at least 3 of the 6 mice included on the whole proteome dataset, we used a random number generator to pick 10 peptides out of the whole proteome array dataset of 6,090,593 unique peptide sequences. The log-transformed fluorescence intensity values associated with these 10 random peptides and the negative control Gria4 peptide, used previously from the whole proteome peptide array, are shown in **Figure 8A**. All 10 of these random peptides showed virtually no reactivity with any of the sera from the 6 immune mice tested, except for one peptide that showed low, but detectible, reactivity with the B2 immune sample on the original whole proteome dataset (Whrn). This one somewhat positive reaction out of the 60 possible combinations of 10 random peptides with 6 serum samples in Figure 8A corresponds to 1.7% positive. We used these 10 random peptides to probe the immune serum samples from the same 20 validation set mice utilized in **Figure 7C** for antibody binding to any of these randomly selected peptides (**Figure 8B**). We observed moderate antibody binding by one of the 20 validation set immune samples (V16) to one of the 10 tested random peptides (Podnl1). No other validation serum samples showed detectible binding to any of these 10 peptides. Thus, of 200 possible combinations of the 20 serum samples with the 10 randomly selected peptides in Figure 8B, only one (0.5%) was positive. Contrasting the relatively absent reactivity of these 20 new validation immune samples to these randomly selected peptides, we now show the relatively strong reactivity of these same 20 validation immune serum samples to the 12 peptides from Figure 7C selected utilizing HERON analyses of the original Nimble data, that were recognized by at least 3 of the original 6 immune mice. These data are shown in **Figure 8C**, using a selection of data also shown in **Figure 7C**. Unlike the 1 positive reaction out of 200 possible combinations shown in Figure 8B, these same 20 immune serum samples now show 20% positive reactions with these 12 HERON selected peptides (48 reactions with an OD reading of >=2 out of 240 possible combinations = 20%). This substantial reactivity of these 20 validation sera to these 12 HERON selected peptides is significantly greater (p< 0.001) from the 0.5% reactivity in the randomly selected peptides.

**Figure 8:**
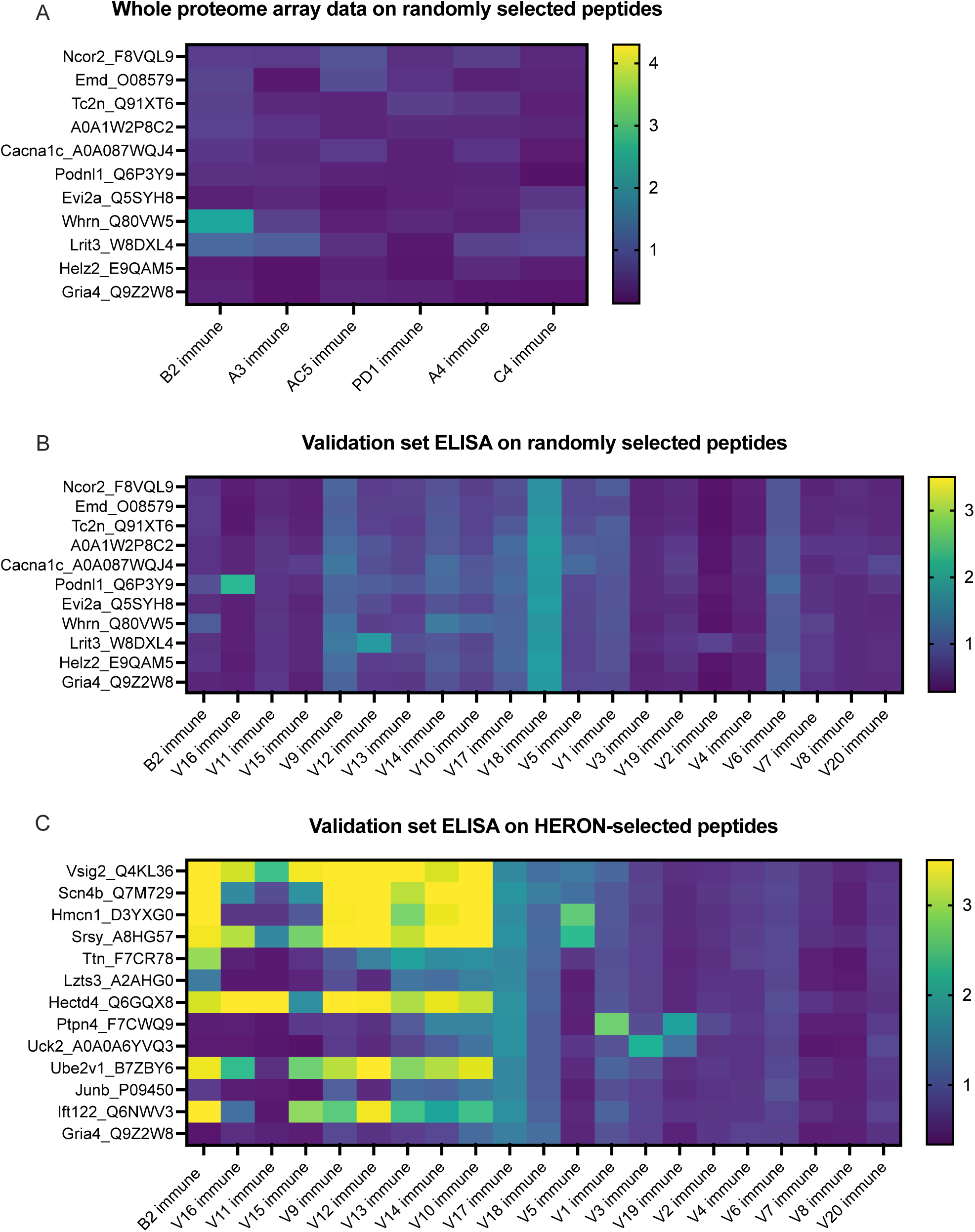
Peptides selected at random from whole proteome array are similarly recognized by ELISA testing for a separate validation set of 20 separate immune mice and show much lower antibody recognition in comparison to HERON-identified peptides present in 50% or more of the original cohort. **A:** Heatmap of 11 peptides from the whole proteome dataset displaying whole proteome peptide array sample results for 6 immune serum samples (the same 6 immune serum samples used for figures 2 & 3). Ten of these 11 Peptides were chosen at random utilizing a random number generator out of all probed peptides from the whole proteome array. One peptide is included as a negative control peptide that was intentionally selected as a negative control; we have never observed antibody binding to it in any of our original or validation tested samples (Gria4, at the bottom of the list). Data shown are log10 of the fluorescence units of the peptide array signal. **B:** ELISA data for the same peptides as in A but using immune mouse serum samples not tested on the whole proteome array from 20 separate immune mice that have received the same treatment to cure their B78 cancer. These 20 new immune serum samples are identical to the 20 new immune serum samples shown in Figure 7C. Also included here is a repeat immune serum sample from one of the 6 immune mice used in the original whole proteome samples as an internal control (B2 immune, also shown in whole proteome data in Figure 8A). Data shown are optical density values read at 450nm length on a scale from 0 to 3.5. **C:** ELISA data for 13 of the 16 peptides highlighted in Figure 7 on the same immune serum samples as in Figure 7C. The peptides included here reflect 12 peptides chosen for strong antibody reactivity in 3 or more of the original 6 mice tested on the whole proteome array. As in Fig. 8A and B, the Gria4 peptide is included as a negative control peptide (at the bottom of the list). Also included here is a repeat immune serum sample from one of the 6 immune mice used in the original whole proteome samples as an internal control (B2 immune, also shown in the whole proteome data in Figure 7A, and ELISA data for original whole proteome samples in Figure 7B). Data shown are optical density values read at 450nm length on a scale from 0 to 3.5. Scales used for the heatmaps in Figure 8 A-C are consistent with the scales used in Figure 7 A-C.

We acknowledge that our sample size of 10 randomly selected peptides in Figure 8B is a small fraction of the over 6 million peptides present on the array. To approach and analyze the ability of the HERON method to identify peptides from the initial Nimble data with the original 6 mice that will show greater than chance reactivity with a new set of immune serum samples, using a calculation that includes a larger number of randomly selected peptides, we employed a model utilizing the hypergeometric distribution (Supplemental Figure 4). When calculating the probability of having a quarter of the previously untested mice recognize a specific peptide with a high ELISA threshold (minimum O.D. signal of 2) if the peptides would have been chosen at random using a hypergeometric test (P(X >= 8 given 14 draws out of a pool of ∼6 million), the chance of having this occur with a separate set of mice is almost zero (**Supplemental Figure 4).** For example, if we assume that 1% (60,906) of the peptides from the set of the ∼6 million unique peptides are reactive, the probability of randomly choosing 14 peptides from the pool of possible peptides and finding that at least 8 of the peptides that are responsive to 25% or more of the mice in the new validation set is 1.9×10^-15^. If we are sampling with replacement from the pool of samples, i.e. using a binomial distribution rather than hypergeometric, the probability is still 2.85×10^-13^. The similar result between the hypergeometric and binomial approach is due to the low likelihood of randomly choosing the same peptide twice amongst the large pool of possible peptides. These analyses indicate that peptides recognized at the moderate level using the Nimble array data for immune sera from 50% of multiple mice are highly likely to be recognized by separate, similarly immunized mice, in a validation set, using ELISA data as a validation system.

Since only 0.45% of peptides tested are recognized at the moderate level by at least one mouse (27639 peptides), and only 0.005% of peptides tested are recognized at the moderate level by 3 or more mice (316 peptides), the fact that 13 of 20 (65%) of independent immune mice from the validation set are recognizing at least one of the 14 peptides (selected from the 0.005% of peptides recognized in the Nimble system by 3 or more mice) by the ELISA system indicates that these peptides co-recognized by multiple mice in the Nimble system are identifying peptides likely to be recognized by independent (validation set) immune mice.

## 5 Discussion

The aim of this study was to establish a method to utilize a high-density overlapping stacked array of peptides representing the entire C57BL/6 proteome in order to identify the “immunome” of epitopes recognized by antibody induced in mice that received curative immunotherapy associated with complete and durable eradication of B78 melanoma tumors with induction of tumor-specific immune memory. In this work, we demonstrated the utility of high-density peptide microarrays for profiling the antibody repertoire in immune serum samples by using a proteome-scale peptide microarray representing all proteins in the mouse proteome. This enabled a fine-mapping of all regions of linear epitopes recognized by circulating antibodies induced during the growth and subsequent complete rejection of a syngeneic murine melanoma. Although these whole-proteome peptide microarrays contained peptides representing the proteome, this study is not a complete analysis of the antibody detected “immunome”. The length of the 16-mer peptides is a limitation, as conformational (or discontinuous) epitopes may remain undetected. Nevertheless, these proteome-scale peptide microarrays, along with the development and use of HERON analytic methods, provided an in-depth snapshot of the information stored in the antibody repertoire of mice immune to B78 melanoma after successful RT+IC immunotherapy.

We were able to achieve improved reliability and reproducibility when considering epitopes rather than single peptide probes (**Figures 2B vs. 2C**). A couple of factors contribute to this; first, there are many more probes than epitopes in the proteome, giving a larger number of possible mismatches. Second, an individual epitope can be a component of several overlapping probes; our HERON algorithm for detecting epitopes recognized by separate assessments of serum samples, requires a degree of similar recognition of the related epitope containing probes by the 2 samples, but does not require complete identity of probe recognition and signal. This enables higher reproducibility of epitopes recognized with high signals than peptides recognized with high signals when replicate chips are evaluated for separate aliquots of the same immune serum sample (**Figures 2B vs. 2C**). Somewhat similarly, when evaluating proteins that are recognized, since a single protein might be recognized by different mice at different regions, the number of proteins recognized by 4, 5 or 6 of the 6 immune mice (at inclusive, moderate and restrictive recognition levels) is substantially higher than the number of epitopes mutually recognized by 4, 5 or 6 of the 6 immune mice (comparing **Figures 3C vs. 4C**).

While we were able to use signal strength as a predictor for peptide binding reliability, it cannot be used as a measure of antibody affinity (Buus *et al*., 2012). Signal strength in peptide arrays is determined by many factors, including quality of synthesized peptide, variations in peptide solvation, presence, or absence of high-affinity antibodies as well as presence or absence of multiple lower-affinity antibodies towards the peptide. As seen in **Figure 3B**, the number of recognized epitopes is similar across all 6 immune mice, while the epitopes recognized by individual mice show a large heterogeneity between mice. This heterogeneity in epitopes recognized is demonstrated by the very large number of epitopes recognized by at least one mouse, compared to the substantially smaller number of epitopes with mutual recognition by any 2 of the 6 immune mice (**Figure 3C**) and only a much smaller fraction of epitopes mutually recognized by 3 or more (50%) of the 6 immune mice. A large heterogeneity in antibody repertoire between individuals has been shown before in humans (Ayoglu *et al*., 2013; Nagele *et al*., 2013) and was expected due to the stochastic nature of V-D-J recombination leading to the specific binding characteristics of an individual antibody generated by a clonally expanded mature B cell.

Interestingly, when validating just a small cohort of 12 peptides, representing ∼2.86% of the peptides examined out of the total of 420 peptides, which were each recognized by at least 3 of our original 6 mice based on the Nimble system data, we were able to show reactivity to at least one of these peptides in 65% of our validation cohort of 20 separate immune mice (**Figure 7D**). While we did not achieve the same rate of recognition for each individual peptide, having at least one peptide recognized by some of these additional 20 mice supports the biological relevance of these proteins being antibody targets by multiple mice in our system. This biological importance is further supported by the testing of random peptides with immune serum samples from 20 additional mice (**Figure 8**) where 10 randomly selected peptides showed only one of the 20 mice recognized just one of the 10 peptides barely above the threshold of an OD value of 2 (mean value of 2.28), corresponding to 0.05% positive reactions of 200). In contrast when these same 20 validation immune serum samples were used to recognize the 12 HERON-selected peptides that showed reactivity with at least 3 of the original 6 mice in the Nimble data, 48 out of the 240 possible combinations had an OD reading of 2 or higher (20%, p< 0.001). This validation indicates that the HERON method of selecting peptides from the Nimble data successfully identifies peptides that are being recognized reproducibly in validation assays at a rate far greater than would be seen merely by chance. More importantly, because the antibody repertoire is determined by stochastic gene rearrangements of V-D-J immunoglobulin gene components, the antibody repertoires of distinct genetically identical mice, should have substantial differences. Thus the ability of the HERON method to identify peptides based on their recognition by an initial set of mice using the Nimble data and these same peptides are subsequently strongly recognized using an independent ELISA assay, on a separate set of previously untested validation immune serum samples, indicates that the peptides (and epitopes) identified by the HERON-method have immunologic importance for other mice from the same strain immunized to the same B78 tumor using the same immunotherapy regimen.

There are several limitations to the current assay configuration to evaluate peptide binding by serum antibodies using a high-density peptide array technology. The assay is set up to provide end-point binding of a complex mixture of antibodies at a single serum dilution. It is difficult to estimate the absolute binding affinity of each antibody clone in the complex sera from such a mixture model. By using different dilutions of known concentrations of well characterized mAbs known to recognize specific epitopes or peptides on the Nimble array, a “standard curve” could be created enabling one to interpolate signals seen with immune sera to the standard curve with the mAb dilutions, allowing calculation of a “mAb concentration equivalent”. We have not pursued this and do not think this has been done yet by others using this technology. By evaluating serial dilutions of multiple different mAbs, at the same concentrations, on this high-density proteome array, one might be able to investigate some general patterns to allow quantitative assessments of binding to elements of the proteome with this technology. Once this knowledge has been acquired, this peptide array might be an optimal way to characterize the binding and specificity/cross-reactivity for new mAbs being developed.

It is also possible that this array misses the antibody target of some clinically important antibody responses. These include antibody reactivity against conformationally determined epitopes that are not generated in relatively small 16-mer peptides. Second, as the peptides used in this array are strictly 16-mer aa sequences, no glycosylation is applied to these peptides. Thus, circulating antibodies that recognize differentially glycosylated peptides would not be detected. Third, some clinically important antibody targets have no peptide component. A major example is the GD2 disialoganglioside, proven to be a clinically important target on neuroblastoma; this glycolipid has no peptide component and is recognized by the Dinutuximab mAb (Yu et al., 2010), and the hu14.18-IL2 immunocytokine used to cure mice of B78 melanoma in this study (Baniel et al., 2020; Morris et al., 2016), and also recognized by circulating antibody in patients immunized with a GD2-containing vaccine (Cheung et al., 2021). This peptide array would not be able to identify antibodies that might have been turned on to such non-peptide antigens, even if they were strongly induced in the process of these mice rejecting, and developing, a memory immune response to these B78 tumors. However, the array would be able to detect antibody binding to peptide mimotopes of such antigens as they are cross-reactive with the non-peptide antigens (Bolesta et al., 2005; Horwacik et al., 2015; Wondimu et al., 2008). Finally, some of the antigenic targets on tumors that have been recognized by adaptive immunity are mutation-driven neo-antigens, with aa sequences different from that controlled by the inherited germline genome. As each individual tumor will have its own unique set of neoantigens, the detection of antibodies to these neoantigens using this high-density peptide array technology would require independently created proteome arrays to be established for each individual tumor being evaluated. This would seem currently impractical.

As such, while other epitope discovery methods are superior in probing more limited numbers of targets to define discontinuous or conformational or glycosylated, or non-peptide, or mutated epitopes and immunodominant responses, this technology appears useful in identifying immunoreactive regions within the entire proteome, not previously considered as potential epitopes, on a large scale.

Beyond the possible utility of identifying biomarkers for effective immune responses induced by cancer immunotherapy, we are hopeful that this technology can be used in profiling antibody responses to many other diseases. This tool was able to detect known and previously unknown protein targets of antibody responses throughout the mouse proteome. This approach could potentially be applied to other cancers to advance diagnostic and cancer vaccine development.

This initial description of the results of our analyses to probe the detection of linear epitopes recognized by sera of mice cured of their B78 tumors, relies on the novel bioinformatic approaches developed to analyze these large data sets, reported separately (McIlwain et al., 2023). This report presents: 1) the immunologic methods used to obtain data and validate it using additional JPT and ELISA systems; 2) the spectrum of peptides, epitopes and proteins recognized; and 3) initial description of what fraction of targets recognized by at least one immune mouse are also recognized by some other mice, despite the stochastic nature of each mouse’s individual B-cell repertoires. Important additional analyses are still underway and are beyond the scope of this initial report. These include characterizing which antigens are recognized by these immune sera and determining their relationship to the B78 melanoma tumor that responded to the immunotherapy in the process of turning on these adaptive antibody responses. These ongoing studies also include identifying which of the proteins recognized by these immune antisera are expressed or over-expressed by the tumor cells themselves, and if expressed by the tumor cells, what is their cellular location (membrane, cytoplasmic or nuclear). Furthermore, even though these antibodies were not seen in naïve mice, and were thus induced by bearing the tumor, and responding to the immunotherapy (as demonstrated in **Figure 5D**), given the very large number of proteins recognized by these immune sera (∼10,000 proteins recognized by at least 1 immune mouse, as shown in **Figure 4C**), it seems very unlikely that all of these are selectively expressed by the tumor cells and not normal tissues. As such these antibodies induced in these mice by implanting and successfully treating these tumors may also reflect antibodies that can recognize proteins from normal tissues and may thereby be considered “auto-antibodies”. Such auto-antibodies may be the mechanism behind auto-immune, paraneoplastic, syndromes seen frequently in patients with cancer (Ong et al., 2022; Villagran-Garcia et al., 2023). Finally, even though multiple distinct proteins are recognized by these immune sera, might some of these antibodies be recognizing shared or similar amino acid sequences on these distinct proteins, reflecting possible immune cross-reactivity of similar antibodies to seemingly distinct proteins? These issues are now being pursued and will be presented in a subsequent separate report (Hoefges et al., 2022).

In summary, this work shows that peptide array technology can be used to detect the linear antibody-recognized “immunome” of sera from mice immune to B78 tumors through RT+IC treatment. While we saw a large heterogeneity between individual mice, some proteins were strongly recognized by sera from multiple immune mice and may potentially be of importance in achieving immunity to the cancer, or as a biomarker of a potent adaptive response to the cancer. This same type of workflow could be applied to other types of cancer or diseases as well as to the analyses of patients that have received effective immunotherapy associated with a clear immune mediated anti-tumor response to their cancer to evaluate the equivalent antibody-recognized human tumor “immunome”. Some work in this cancer realm, and in analyses of auto-immunity and anti-viral immunity has been reported and is underway (Heffron et al., 2021; Mergaert et al., 2022; Potluri et al., 2020; Zheng et al., 2021).

## 8 Ethics Statement

The animal study was reviewed and approved by Institutional Animal Care and Use Committee, University of Wisconsin-Madison.

## 9 Conflict of Interest

RSP, BG & JP are all employees of Nimble Therapeutics, the producer of the high-density peptide arrays used for this research. Other than these affiliations, the authors declare that the research was conducted in the absence of any commercial or financial relationships that could be construed as a potential conflict of interest.

## 10 Author Contributions

AH designed the experiments, collected the data, designed, and performed the analysis, interpreted the data, generated the figures, and drafted the manuscript for this study. SJM designed and performed the analysis, generated figures, and assisted in writing and editing the manuscript. NM, AX & DM assisted in data collection and analysis and editing the manuscript. KT, TL and KK helped in the analysis of experiments and editing the manuscript. RSP, BG & JP assisted in design and execution of experiments and editing of the manuscript. MH, AF & NT assisted in collecting the data and editing the manuscript. JAH and ZSM assisted in designing the experiments and editing the manuscript. AKE assisted in designing the experiments and interpreting the data as well as editing the manuscript. PMS and IMO assisted in designing the experiments and analysis, interpreting the data, and editing the manuscript. All authors contributed to the article and approved the submitted version.

## 11 Funding

This work was supported by Midwest Athletes Against Childhood Cancer; Stand Up 2 Cancer and CRUK; the St. Baldrick’s Foundation; the Crawdaddy Foundation; the University of Wisconsin Carbone Cancer Center; the Data Science Initiative grant from the University of Wisconsin Office of the Chancellor and the Vice Chancellor for Research and Graduate Education, The Cancer Research Institute, Alex’s Lemonade Stand Foundation, and the Children’s Neuroblastoma Cancer Foundation. This research was also supported in part by public health service grants U54-CA232568, R35-CA197078, U01-CA233102, and Project 3 of P01CA250972 from the National Cancer Institute; Clinical and Translational Science Award (CTSA) program (ncats.nih.gov/ctsa), through the National Institutes of Health National Center for Advancing Translational Sciences (NCATS), grants UL1TR002373 and KL2TR002374; 2U19AI104317-06U19-2 from the National Institute of Allergy and Infectious Diseases; and UL1TR002373 from the National Institutes of Health and the Department of Health and Human Services. Shared resource flow cytometry reagents and equipment are funded by University of Wisconsin Carbone Cancer Center Support Grant P30 CA014520. The content is solely the responsibility of the authors and does not necessarily represent the official views of the National Institutes of Health.

## 7 List of abbreviations

GD2: Disialoganglioside
IC: immunocytokine
ICC: Intraclass correlation coefficient
IT: intratumoral
ISV: *in situ* vaccine
O.D.: optical density
RT: radiation

## 12 Acknowledgments

We would like to thank Drs. Alexander Rakhmilevich, Lauren Zebertavage, Ravi Patel, and Claire Baniel, as well as Alina Hampton and Clinton Heinze for helpful conversations and assistance in sample collection. We would also like to thank the University of Wisconsin Carbone Cancer Center Flow Cytometry Laboratory for facilities and services. The schematic representations in Figure 1 were created with BioRender.com

## Footnotes

1 To simplify our explanation, we use the binomial distribution, which assumes the probability is the same for every trial (assumes replacement): if there are 10000 positive peptides within the pool of total possible peptides (6000000), the probability of getting k=8 positive peptides out of n=14 selected peptides (if all peptides are equally likely) is p=((6000000 – 10000)/6000000)^k^. The probability of 14-k failures is (1-p)^(14-k)^. However, there are 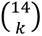 different ways of distributing k=8 successes in a sequence of n=14 trials, so 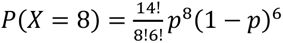. Finally, we calculate the sum of the probabilities using 8 to 14 peptides to obtain the cumulative distribution function P(X >= 8).

## Notes

https://doi.org/10.5281/zenodo.7871565

### Summary of Updates

Figure 8 added, we acquired additional data important for this manuscript

