## Supplemental Figures for "Antibody landscape of C57BL/6 mice cured of B78 melanoma via immunotherapy"

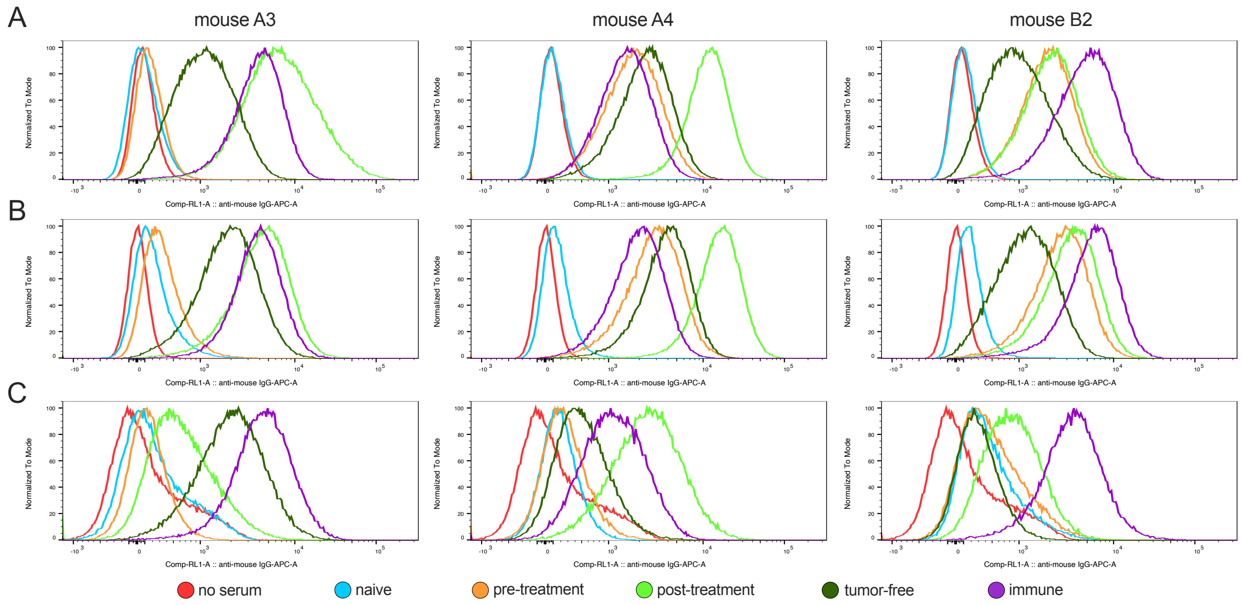


**Supplemental Figure 1:** Histograms of IgG binding to tumor cells for three individual mice (A3, A4 and B2) on 3 different tumor cell lines.

Timepoints correspond to the sample collection timeline in Figure 1A. **A:** Binding of serum antibodies to B16 tumor cells as measured via flow cytometry. **B:** Binding of serum antibodies to B78 tumor cells as measured via flow cytometry. **C:** Binding of serum antibodies to Panc02 tumor cells as measured via flow cytometry. Data are shown as fluorescence intensities detected in the red channel measuring fluorescence signal for APC. The samples for each individual mouse are normalized to mode to enable comparison between the different time points for each mouse.

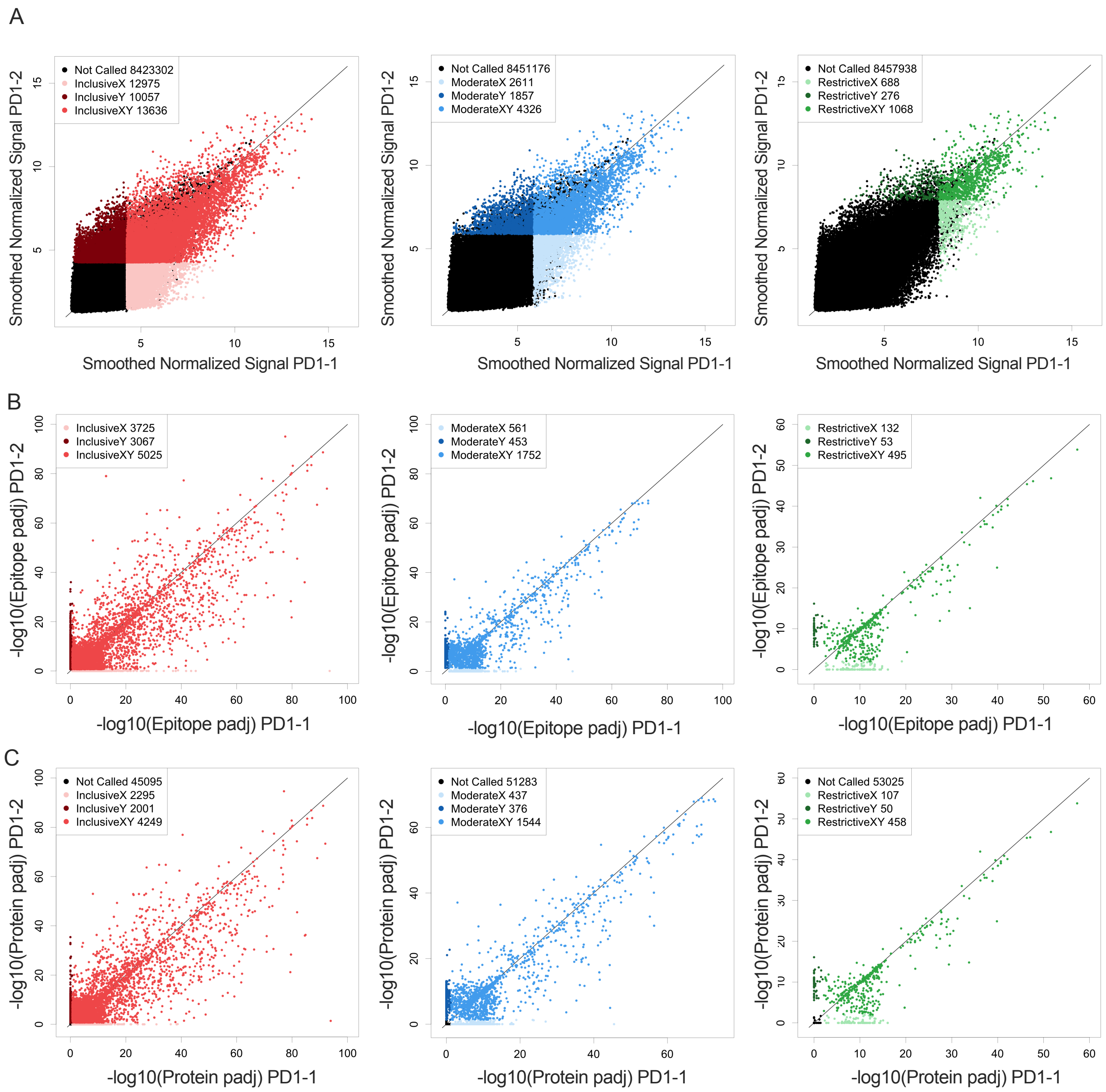


**Supplemental Figure 2:** Reproducibility and reliability of probe, epitope, and protein calls

**A-C**: Reliability of Peptide array: Called peptides were separated in 3 categories [left panel inclusive; middle panel moderate; right panel restrictive] based on signal strength of the bound peptides. **A:** Cryopreserved serum samples from the same Immune blood sample from mouse PD1, were tested independently (sample PD1-1 and PD1-2) on identical whole proteome chips, in assays that were performed 1 year apart. These showed high correlation between signal strength and repeatability of results after data pre-processing on a probe level. Shown are the log transformed processed raw peptide array data generated in arbitrary fluorescence units. In the inclusive category (signal >3xSD greater than the mean) 58% of probes recognized in sample PD1-2 are also recognized in sample PD1-1. In the moderate category (signal >6xSD greater than the mean) 70% of probes recognized in sample PD1-2 are also recognized in sample PD1-1 and in the restrictive category (Signal > 10xSD greater than the mean) 80% of probes recognized in sample PD1-2 are also recognized in sample PD1-1. Shown are the log transformed values of background corrected smoothed raw peptide array data generated in arbitrary fluorescence units. **B:** Scatter plot of the same serum sample from **A** looking at epitope level data instead of individual peptides. Plotted are the -log10 values of the epitope p values for each epitope recognized in sample PD1-1 and sample PD1-2. Graphs are split up into each category, B-left, showing inclusive category with 43% of epitopes co-recognized by both samples, B-middle showing moderate category with 63% of epitopes co-recognized by both samples and B-right showing restrictive category with 73%. of epitopes co-recognized by both samples, **C:** Scatter plot of the same serum sample from A & B looking at protein level data instead of individual peptides or epitopes. Plotted are the -log10 values of the protein p values for each protein recognized in sample PD1-1 and sample PD1-2. Graphs are split up into each category, C-left showing inclusive category with 50% of proteins co-recognized by both samples, C-middle showing moderate category with 66% of proteins co-recognized by both samples and C-right showing restrictive category with 75% of proteins co-recognized by both samples.

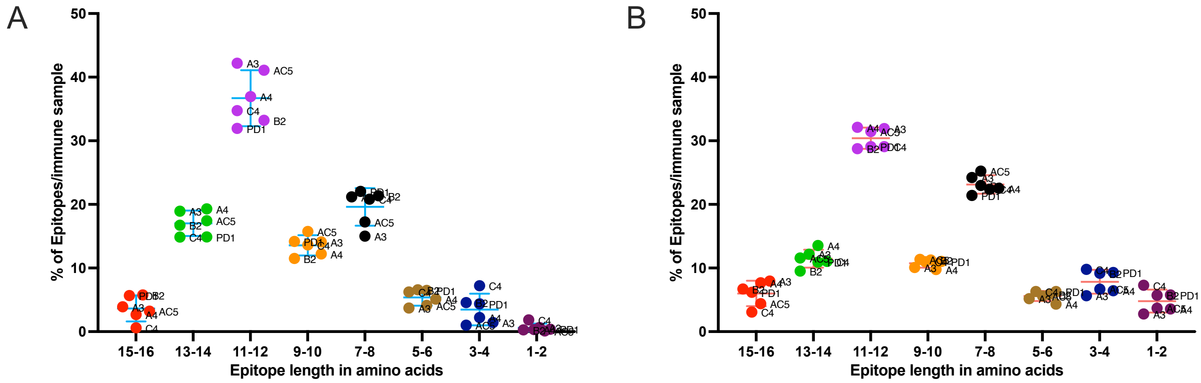


**Supplemental Figure 3:** Epitope length divided by intensity of binding signal

**A & B**: Epitopes were split in epitope length spanning between 1 and 8 consecutive 16-mer probes tiled at 2 amino acids. Over 80% of epitopes required amino acid sequences of 7-16 amino acids for binding in the moderate (**A**) and inclusive category(**B**). Only linear epitopes were probed, but small conformational epitopes are possible within a 16-mer peptide. The X axis shows the length of the detected epitope in 2aa steps as most peptides were tiled at 2 amino acids. The Y-axis shows the percentage of all epitopes per sample, for each of the 6 mice tested, with the corresponding epitope length. The designation for each of the individual 6 mice is shown in small letters next to each of the 6 dots appearing in each of the columns.

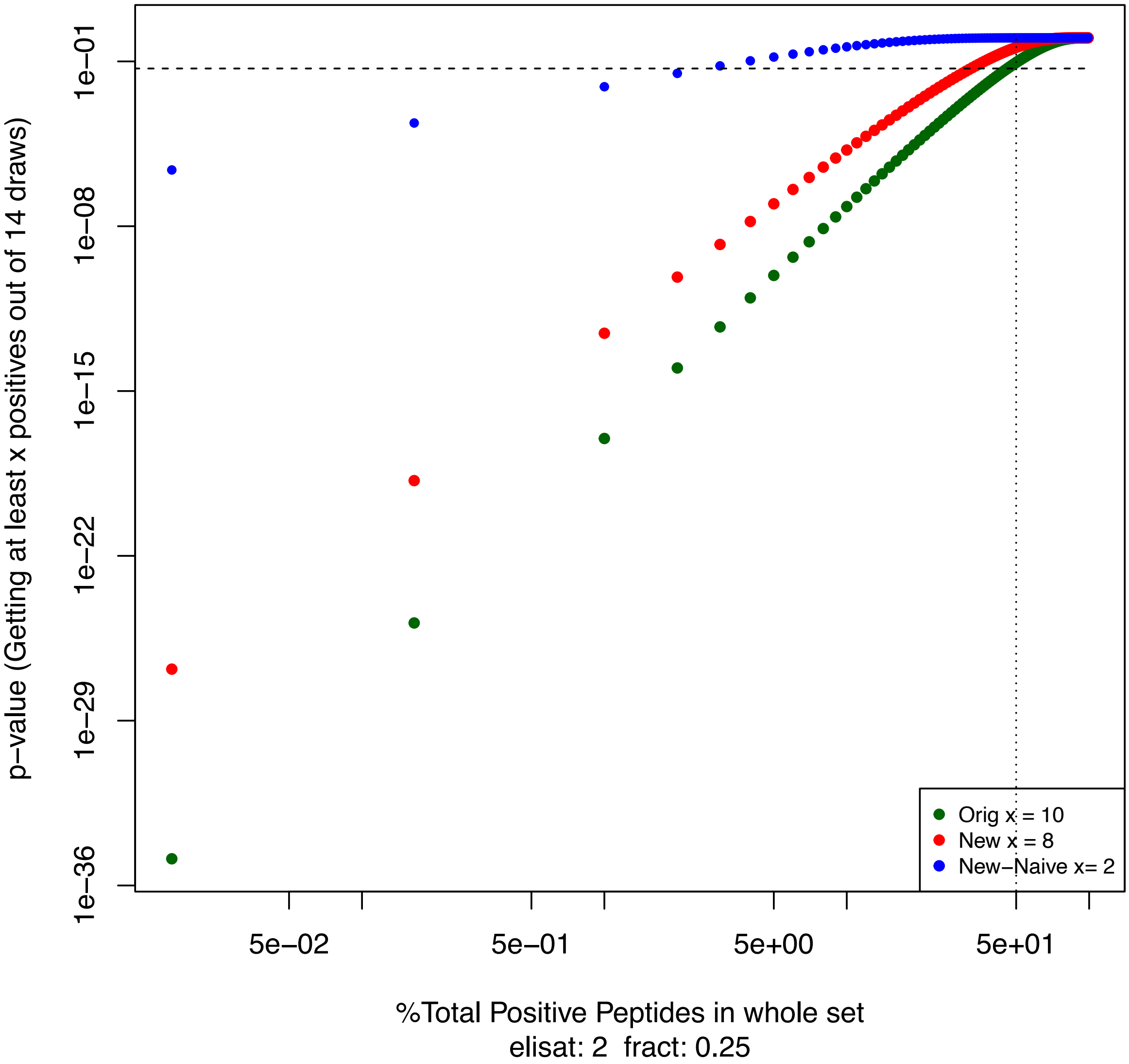


**Supplemental Figure 4:** Hypergeometric distribution of proposed positive peptides. Plot of the hypergeometric p-value of getting at least *x* or more positives out of a sample of 14 randomly chosen peptides versus the % of proposed total peptides in the whole set of ~6.5 million unique peptides. Green - original ELISA results on immune mice, Red - validation ELISA results on immune mice, Blue - validation ELISA results on Naive mice. Both the red and green line show very significant p-values for up to almost 50% of peptides of the 6.5million unique peptides giving a positive signal. However, looking at the Naïve samples a significant p-Value is only reached by a maximum of 5% of positive peptides in this simulation using values derived from the Nimble and ELISA data generated.

**Supplemental Table 1:** List of serum samples used for the separate assays using whole proteome peptide arrays, JPT multi-well peptide array and peptide ELISA, with data displayed in this manuscript. Identifiers refer to individual mouse IDs, used for naïve as well as immune samples (noted in the column labeled "status"). Naïve pool 1, naïve pool 2, immune pool 1, and immune pool 2 are pooled samples consistent of 4 individual mouse samples each with values only used and displayed in the boxplot in Figure 2a. All other figures do not contain pooled sample values. Naïve pool 1 consists of individual mouse IDs PD1, PD4, PD5 and AC5. Naïve pool 2 consists of individual mouse IDs AC5, AC1, AB4 and PD1. Immune pool 1 consists of individual mouse IDs PD1, PD4, AB3 and AC5. Immune pool 2 consists of individual mouse IDs AC5, CS4, PD1 and CS6. These pooled samples were used at a 1:50 dilution of serum which resulted in an individual sample serum concentration of 1:200. The pooled samples were omitted for the rest of the paper as it was not clear which bound peptides resulted from which serum sample tested and did not allow for individual mouse to mouse comparisons.

| Identifier | Status | Nimble whole proteome array 1 | Nimble whole proteome array 2 | JPT multiwell peptide array | ELISA |
|---|---|---|---|---|---|
| AC5 | naïve | X | | | |
| | immune | X | | X | X |
| PD1 | naïve | X | | | X |
| | immune | X | X | X | X |
| B2 | naïve | | X | X | X |
| | immune | | X | X | X |
| A3 | naïve | | X | X | X |
| | immune | | X | X | X |
| C4 | immune | | X | X | |
| A4 | naïve | | X | | X |
| | immune | | X | X | X |
| naïve pool 1 | naive | X | | | |
| naïve pool 2 | naive | X | | | |
| immune pool 1 | immune | X | | | |
| immune pool 2 | immune | X | | | |
| Validation cohort for ELISA only: | | | | | |
| V1 | naïve | | | | X |
| | immune | | | | X |
| V2 | naïve | | | | X |
| | immune | | | | X |
| V3 | naïve | | | | X |
| | immune | | | | X |
| V4 | naïve | | | | X |
| | immune | | | | X |
| V5 | naïve | | | | X |
| | immune | | | | X |
| V6 | naïve | | | | X |
| | immune | | | | X |
| V7 | naïve | | | | X |
| | immune | | | | X |
| V8 | naïve | | | | X |
| | immune | | | | X |
| V9 | naïve | | | | X |
| | immune | | | | X |
| V10 | naïve | | | | X |
| | immune | | | | X |
| V11 | naïve | | | | X |
| | immune | | | | X |
| V12 | naïve | | | | X |
| | immune | | | | X |
| V13 | naïve | | | | X |
| | immune | | | | X |
| V14 | naïve | | | | X |
| | immune | | | | X |
| V15 | immune | | | | X |
| V16 | immune | | | | X |
| V17 | immune | | | | X |
| V18 | immune | | | | X |
| V19 | immune | | | | X |
| V20 | immune | | | | X |

**Supplemental Table 2:**

Values corresponding to Figure 2 and supplemental Figure 2 and percentage calculations based on the respective called probes, epitopes, or proteins for serum samples from the same mice at the same timepoints (using the same cut-offs as shown in Fig. 2 and Suppl. Fig. 2) either run within a day of each other (for mouse B2 in Fig. 2) or a year apart (for mouse PD1, in Supplemental Figure 2). In addition: calculations comparing serum from 2 different mice tested in the same run (B2 vs repeat of PD1) show that different mice demonstrate co-recognition of a small fraction of the samples seen by those same individual mice. Calculations for % were done as following: overall=X only +Y only + X&Y; % overall=(overall/all probes)x100; % X&Y=(X&Y/all probes)x100; % of called (X)=(X&Y/(X only + X&Y))/100; % of called (Y)=(X&Y/(Y only + X&Y))x100; % of called X&Y=(X&Y/overall)x100.

| | Sample | X (.1) only | Y (.2) only | X&Y | overall | all probes | not called | % overall | % X&Y | % of called (X) | % of called (Y) | % of called X&Y |
|---|---|---|---|---|---|---|---|---|---|---|---|---|
| **Probe** | B2 Z3 | 5311 | 7178 | 22806 | 35295 | 8459970 | 8424675 | 0.417 | 0.270 | 81.11 | 76.06 | 64.62 |
| | PD1 Z3 | 12975 | 10057 | 13636 | 36668 | 8459970 | 8423302 | 0.433 | 0.161 | 51.24 | 57.55 | 37.19 |
| | B2 Z6 | 591 | 1027 | 6653 | 8271 | 8459970 | 8451699 | 0.098 | 0.079 | 91.84 | 86.63 | 80.44 |
| | PD1 Z6 | 2610 | 1857 | 4326 | 8793 | 8459970 | 8451177 | 0.104 | 0.051 | 62.37 | 69.97 | 49.20 |
| | B2 Z10 | 95 | 149 | 1422 | 1666 | 8459970 | 8458304 | 0.020 | 0.017 | 93.74 | 90.52 | 85.35 |
| | PD1 Z10 | 688 | 276 | 1068 | 2032 | 8459970 | 8457938 | 0.024 | 0.013 | 60.82 | 79.46 | 52.56 |
| | B2vsPD1 (same run, different mice) | | | | | | | | | | | |
| | Z3 | 25536 | 21112 | 2581 | 49229 | 8459970 | 8410741 | 0.582 | 0.031 | 9.18 | 10.89 | 5.24 |
| | Z6 | 6408 | 5347 | 836 | 12591 | 8459970 | 8447379 | 0.149 | 0.010 | 11.54 | 13.52 | 6.64 |
| | Z10 | 1355 | 1128 | 216 | 2699 | 8459970 | 8457271 | 0.032 | 0.003 | 13.75 | 16.07 | 8.00 |
| **Epitope** | B2 Z3 | 1374 | 1724 | 7868 | 10966 | | | | | 85.13 | 82.03 | 71.75 |
| | PD1 Z3 | 3725 | 3067 | 5025 | 11817 | | | | | 57.43 | 62.10 | 42.52 |
| | B2 Z6 | 58 | 89 | 2376 | 2523 | | | | | 97.62 | 96.39 | 94.17 |
| | PD1 Z6 | 561 | 453 | 1752 | 2766 | | | | | 75.75 | 79.46 | 63.34 |
| | B2 Z10 | 6 | 2 | 526 | 534 | | | | | 98.87 | 99.62 | 98.50 |
| | PD1 Z10 | 132 | 53 | 495 | 680 | | | | | 78.95 | 90.33 | 72.79 |
| | B2vsPD1 (same run, different mice) | | | | | | | | | | | |
| | Z3 | 8036 | 6886 | 1206 | 16128 | | | | | 13.05 | 14.90 | 7.48 |
| | Z6 | 2434 | 1844 | 361 | 4639 | | | | | 12.92 | 16.37 | 7.78 |
| | Z10 | 427 | 443 | 105 | 975 | | | | | 19.74 | 19.16 | 10.77 |
| **Protein** | B2 Z3 | 806 | 1137 | 6440 | 8383 | 53640 | 45257 | 15.628 | 12.006 | 88.88 | 84.99 | 76.82 |
| | PD1 Z3 | 2295 | 2001 | 4249 | 8545 | 53640 | 45095 | 15.930 | 7.921 | 64.93 | 67.98 | 49.72 |
| | B2 Z6 | 45 | 73 | 2166 | 2284 | 53639 | 51355 | 4.258 | 4.038 | 97.96 | 96.74 | 94.83 |
| | PD1 Z6 | 437 | 375 | 1544 | 2356 | 53639 | 51283 | 4.392 | 2.879 | 77.94 | 80.46 | 65.53 |
| | B2 Z10 | 4 | 3 | 501 | 508 | 53640 | 53132 | 0.947 | 0.934 | 99.21 | 99.40 | 98.62 |
| | PD1 Z10 | 107 | 50 | 458 | 615 | 53640 | 53025 | 1.147 | 0.854 | 81.06 | 90.16 | 74.47 |
| | B2vsPD1 (same run, different mice) | | | | | | | | | | | |
| | Z3 | 5058 | 4062 | 2188 | 11308 | 53640 | 42332 | 21.081 | 4.079 | 30.20 | 35.01 | 19.35 |
| | Z6 | 1767 | 1475 | 445 | 3687 | 53640 | 49953 | 6.874 | 0.830 | 20.12 | 23.18 | 12.07 |
| | Z10 | 404 | 407 | 101 | 912 | 53640 | 52728 | 1.700 | 0.188 | 20.00 | 19.88 | 11.07 |

**Supplemental Table 3:**
284 peptides tested on JPT, and Nimble platform used in Figure 5 in no or high signal category. Data provided for all samples from Nimble and JPT shown in Figure 5. Categories were chosen based on Nimble whole proteome array 1 and then tested on JPT and Nimble whole proteome array 2. All data from JPT multi-well peptide array runs and matched Nimble peptides on all samples can be found in Supplemental Data 1.

**Supplemental Data 1:**
376 peptides tested on JPT and Nimble platform. Data provided for all samples from Nimble and JPT runs performed. Categories were chosen based on Nimble whole proteome array 1 and then tested on JPT and Nimble whole proteome array 2.
